# Functional resilience of mutually repressing motifs embedded in larger networks

**DOI:** 10.1101/2022.01.13.475824

**Authors:** Pradyumna Harlapur, Atchuta Srinivas Duddu, Kishore Hari, Mohit Kumar Jolly

## Abstract

Elucidating the design principles of regulatory networks driving cellular decision-making has important implications in understanding cell differentiation and guiding the design of synthetic circuits. Mutually repressing feedback loops between ‘master regulators’ of cell-fates can exhibit multistable dynamics, thus enabling multiple “single-positive” phenotypes: (high A, low B) and (low A, high B) for a toggle switch, and (high A, low B, low C), (low A, high B, low C) and (low A, low B, high C) for a toggle triad. However, the dynamics of these two network motifs has been interrogated in isolation *in silico*, but *in vitro* and *in vivo*, they often operate while embedded in larger regulatory networks. Here, we embed these network motifs in complex larger networks of varying sizes and connectivity and identify conditions under which these motifs maintain their canonical dynamical behavior, thus identifying hallmarks of their functional resilience. We show that an increased number of incoming edges onto a motif leads to a decay in their canonical stand-alone behaviors, as measured by multiple metrics based on pairwise correlation among nodes, bimodality of individual nodes, and the fraction of “single-positive” states. We also show that this decay can be exacerbated by adding self-inhibition, but not self-activation, loops on the ‘master regulators’. These observations offer insights into the design principles of biological networks containing these motifs, and can help devise optimal strategies for integration of these motifs into larger synthetic networks.

## Introduction

Gene Regulatory Networks (GRNs) are an integral part of the control structure involved in various cellular processes such as cell-fate decisions made during embryonic development, cellular reprogramming, and phenotypic switching among two or more cell types. A pluripotent cell is capable of differentiating to more than one cell type in response to varying stimuli. This property of coexistence of more than one stable steady state (phenotypes) is referred to as multi-stability and it underlies the dynamics of many GRNs involved in decision-making during differentiation ^1^. Such multi-stability has been seen during cellular reprogramming as well as phenotypic switching under many circumstances. Thus, elucidating the dynamical principles of multi-stable GRNs and network motifs holds promise for understanding many biological processes and control applications in synthetic biology^2–5^.

One of the most frequently observed and extensively investigated network motifs is the ‘Toggle Switch’ (TS), i.e., two mutually repressing regulators A and B, each driving a different cell fate ^6– 8^. The TS enables two mutually exclusive “single-positive” outcomes - (high A, low B) and (low A, high B), thus showing bistable dynamics and allowing a pluripotent cell to choose from two cell fates ^2,8,9^. For instance, PU.1 and GATA1 form a TS that drives hematopoietic stem cells to either a common myeloid progenitor (PU.1 high, GATA1 low) or an erythroid one (PU.1 low, GATA1 high) ^2,10^. Also, in *Escherichia coli*, the construction of a TS exhibiting bistability and switching between the two states in response to external signals has driven an extensive design of synthetic genetic circuits ^8,11,12^. Another network motif is a ‘Toggle Triad’ (TT), i.e., three mutually repressing regulators A, B, and C, each driving a respective cell fate ^13–16^. TT can enable a progenitor cell to differentiate into three distinct cell fates; for instance, the case of naïve CD4+ T helper cells differentiating to Th1, Th2, and Th17 cells ^13,17,18^. The three canonical “single-positive” states enabled by TT are: (high A, low B, low C), (low A, high B, low C) and (low A, low B, high C), as also seen in a recent synthetically constructed TT based on protein dimerization instead of transcriptional regulation ^19^.

While the dynamics of TS and TT have been extensively investigated deterministically and stochastically, most such investigations have considered them in isolation, i.e. TS or TT are assumed to be not connected to any other network components ^13–15,20–22^. However, in reality, a TS or TT is only a small part embedded in a larger network of interconnected proteins and signaling components. Here, we investigate and quantify the behavior of TS and TT network motifs when embedded in much larger networks, using three properties: bimodality of individual nodes, pairwise correlation coefficient between nodes, and fraction of canonical “single-positive” states. We noticed that for a TS, an increase in the number of incoming edges on the two nodes of a TS (i.e. in-degrees for A and B) resulted in deviation from stand-alone behavior, as captured by changes in all the three abovementioned properties. Further, an asymmetry in the in-degrees for both nodes also compromised bistability. However, for a TT, the fraction of “single-positive” (F1) states and maximum correlation coefficient (MaxCC) were reliable metrics to quantify its deviation from stand-alone dynamics. We observed that as the net in-degree of a TT increased, “single-positive” steady states (e.g. (high A, low B, low C)) were replaced by “double-positive” ones (e.g. (high A, high B, low C)). The value of the maximum of the three pairwise correlation coefficients (Max CC, i.e. maximum of (CC AB, CC BC, CC AC)) also increased with increasing in-degree of TT. These observations suggest that in addition to the previously studied factors influencing the dynamics of TS or TT, the local density around these motifs, when embedded in larger networks, could also influence their functional properties.

## Results

### Stand-alone properties of toggle switch and toggle triad

We first investigated the stand-alone properties of a toggle switch (TS) and toggle triad (TT), before embedding them in larger networks. RACIPE formalism ^23^ was used to simulate these networks for 10,000 randomized parameter sets sampled from a predetermined parameter space. Three such replicates (each with 10,000 parameter sets * 100 initial conditions per parameter set) were performed for each motif. The resultant steady state values for each node were normalized and converted to z-scores. Pairwise correlation coefficients (CC) were then calculated between the steady state values of nodes in a TS or TT. Also, for the TS, we calculated Sarle’s bimodality coefficient (BiC) ^24^ values for each node. BiC values range from 0 to 1, with values closer to 1 representing higher bimodality, and any value above 0.55 considered to represent a bimodal distribution ^23^.

The TS motif (**Fig 1A**) mainly showed two “single-positive” steady states: ((low A, high B); (A, B) = (0, 1)) and ((high A, low B); (A, B) = (1, 0)), as observed in the bivariate plot (**Fig 1B**). The steady state values of both nodes in a TS showed a bimodal distribution (BiC A = BiC B = 0.78), with the peaks representing the corresponding high and low steady state values. Because the two nodes of a TS repress each other, the correlation coefficient between steady state values of the nodes was strongly negative (CC AB = - 0.83) (**Fig 1C**). The steady state values for a TT motif (**Fig 1D**), for any given pair of TT nodes, had three distinct clusters, which represent the three “single-positive” stable steady states namely ((low A, low B, high C); (A, B, C) = (0, 0, 1)) state, ((low A, high B, low C); (A, B, C) = (0, 1, 0)) state and ((high A, low B, low C); (A, B, C) = (1, 0, 0)) state. One node shows higher expression in these states while the other two nodes have repressed expression (**Fig 1E, S1A-B**). In contrast to TS, the average BiC of a node in TT is 0.43 (standard deviation = 0.004), implying a more unimodal-like distribution with little difference in the high and low steady state values of a node in a TT. Although negative, the magnitude of pairwise correlation coefficient between the steady state values of any two nodes of a TT was less than that of a TS (CC AB = CC AC = - 0.39, CC BC = -0.40) (**Fig 1F, S1C-D**). This decrease could be because although any pair of two nodes mutually repress, due to the dominance of “single-positive” steady states, two nodes of TT can still show similar low-expression steady state values, thus leading to a relatively lower magnitude of correlation coefficient between them. On the other hand, two nodes of a TS are strictly confined to having opposing expression profiles, leading to a strongly negative correlation.

**Figure 1:**
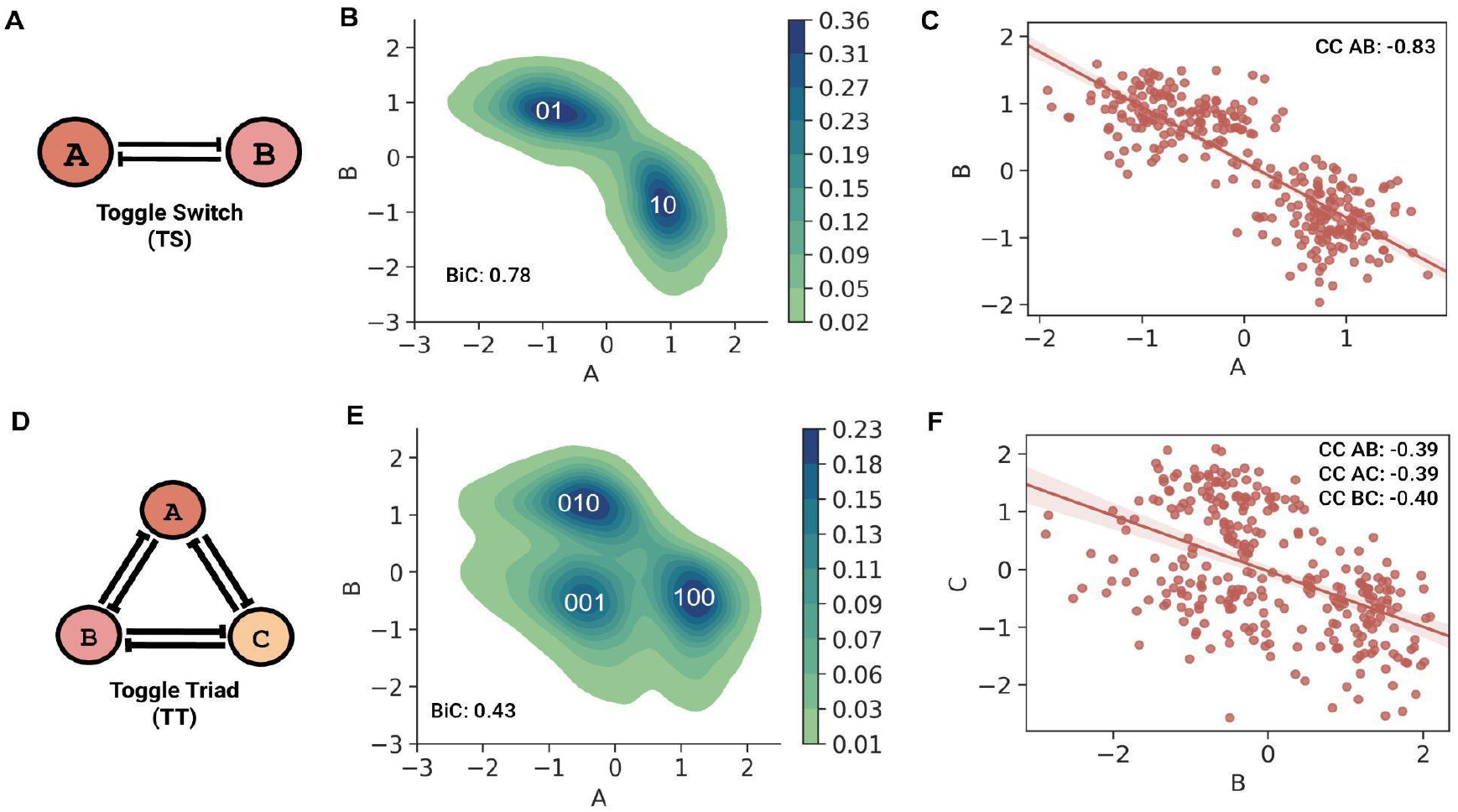
Stand-alone dynamics of two-node and three-node mutually repressing motifs. **A)** A toggle switch (TS) motif comprises two nodes A and B that mutually inhibit each other. **B)** Probability density plot of steady state values of nodes in a TS. The two dense clusters correspond to “single-positive” 01 and 10 steady states (F1) of a TS. **C)** Regression plot between the steady states values of two nodes, A and B of a TS. Correlation coefficient (CC AB) between them is -0.83. **D)** A toggle triad (TT) motif comprises three mutually repressing nodes A, B, and C. **E)** Probability density plot of steady-state values of two TT nodes A and B; the three clusters represent three “single-positive” steady states (F1), 001, 010, and 100. **F)** Regression plot between steady state values of nodes B and C of a TT. Correlation coefficient (CC BC) = -0.39. Other pairwise correlation coefficient values are also mentioned: CC (AC) = -0.39, CC (AB) = -0.40.

### Functional traits of toggle switch depend on density rather than the size of the larger networks it is embedded in

Next, we embedded TS and TT motifs in different larger networks having combinations of four different network orders and three distinct densities (mean connectivity) to understand how the above mentioned stand-alone dynamic traits of TS and TT change. The four network orders are 5N, 10N, 15N and 20N, where N is the number of nodes in a network in which these motifs were embedded. The three mean connectivity values are E:2N, E:4N, and E:6N, where E:xN signifies that the number of edges (E) is x times the number of nodes (N). The combinations of the four network orders (5N, 10N, 15N and 20N) and three mean connectivity (E:2N, E:4N and E:6N) resulted in twelve different types of networks (**Fig 2A**). For each type of network, n=100 random network topologies were generated. TS motifs were then embedded into these 1200 (12 types x n=100) randomly generated networks to study the motif’s behavior. For instance, a TS embedded in a 5N, E:4N network will have 7 nodes (5 nodes + 2 nodes of TS) and 22 edges (5*4 edges + 2 edges of TS) (**Fig 2B**). The same process was repeated for TT motif to generate 1200 random networks. RACIPE was used to simulate dynamics of these larger networks of varying sizes and mean connectivity, with each network being simulated three times **(Fig 2C)**. The generated outputs were normalized by z-scoring and then analyzed to characterize the behavior of TS and TT motifs upon embedding. Three metrics were assessed to quantify the dynamic resilience of TS and TT motifs when embedded in larger networks: bimodality coefficient (BiC), correlation coefficient (CC) and frequency of canonical ‘single-positive’ states as a fraction of all steady states observed (F1).

**Figure 2:**
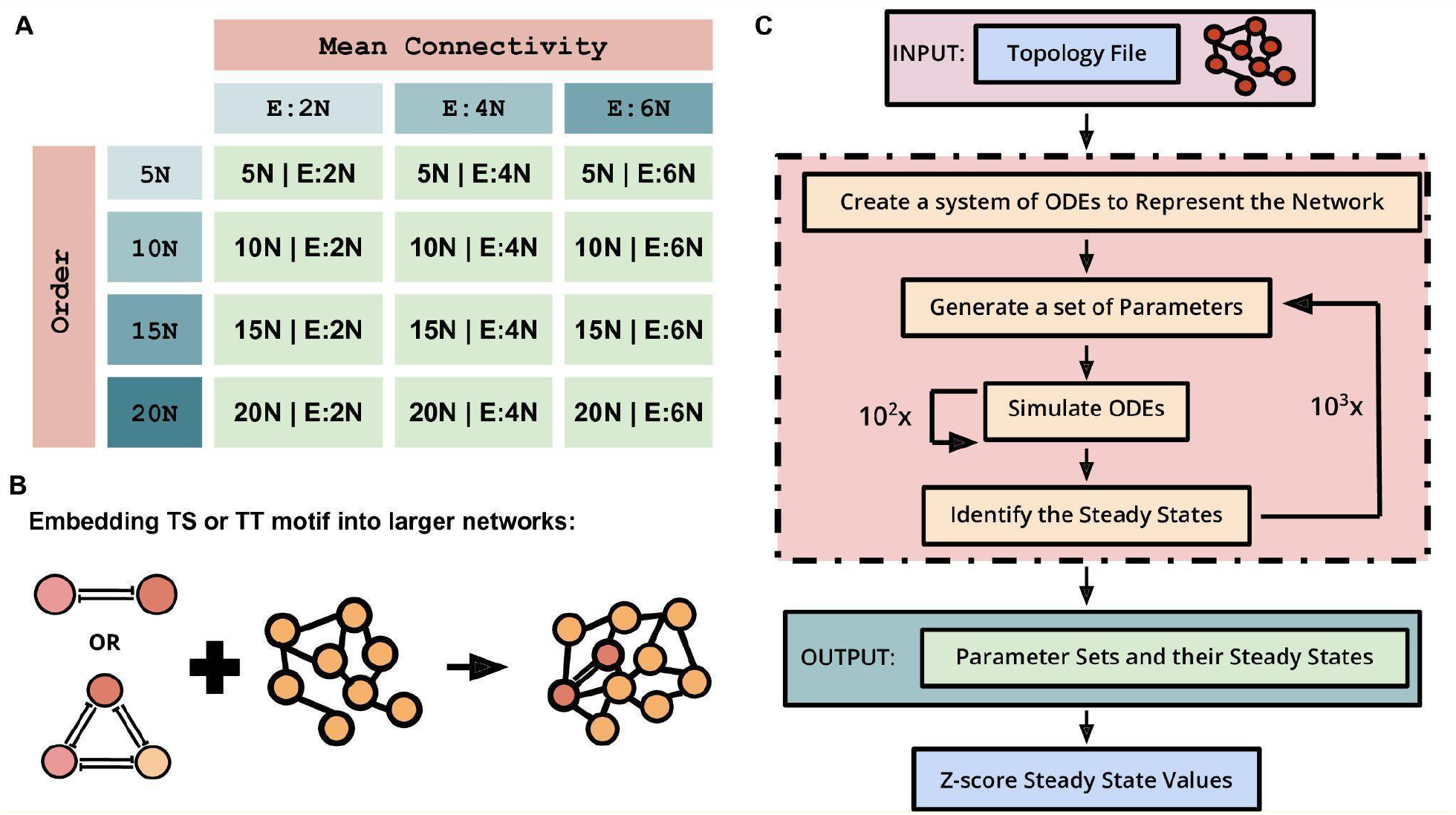
Schematic of the pipeline used to generate, simulate and analyze the motifs embedded in random networks. **A)** Table showing the twelve types of networks which were created as combinations of three mean connectivity and four network orders. **B)** Schematic showing the process of embedding the motifs into the created networks. **C)** Simulation pipeline of RACIPE used to get the steady state values for these larger networks containing embedded TS or TT, for further analysis.

To evaluate how network size and density can independently influence the behavior of a TS when embedded in larger networks of varying orders and mean connectivity, we compared the behavior for networks sharing the same mean connectivity but having different network orders or *vice versa*. Interestingly, for networks with same mean connectivity, CC between the two nodes in a TS (CC AB) did not show any significant variation for varied network orders (**Fig 3A, i**). However, when controlling for network order, CC AB reduced as the mean connectivity increased (**Fig 3A, ii**). A similar trend, i.e., the dependence on network density rather than on network size and a decrease in magnitude with increasing network density, was also observed in distributions of BiC values: BiC A and BiC B **(Fig 3B, i-ii; S1E-F)** and for F1 **(Fig 3C, i-ii)**.

**Figure 3:**
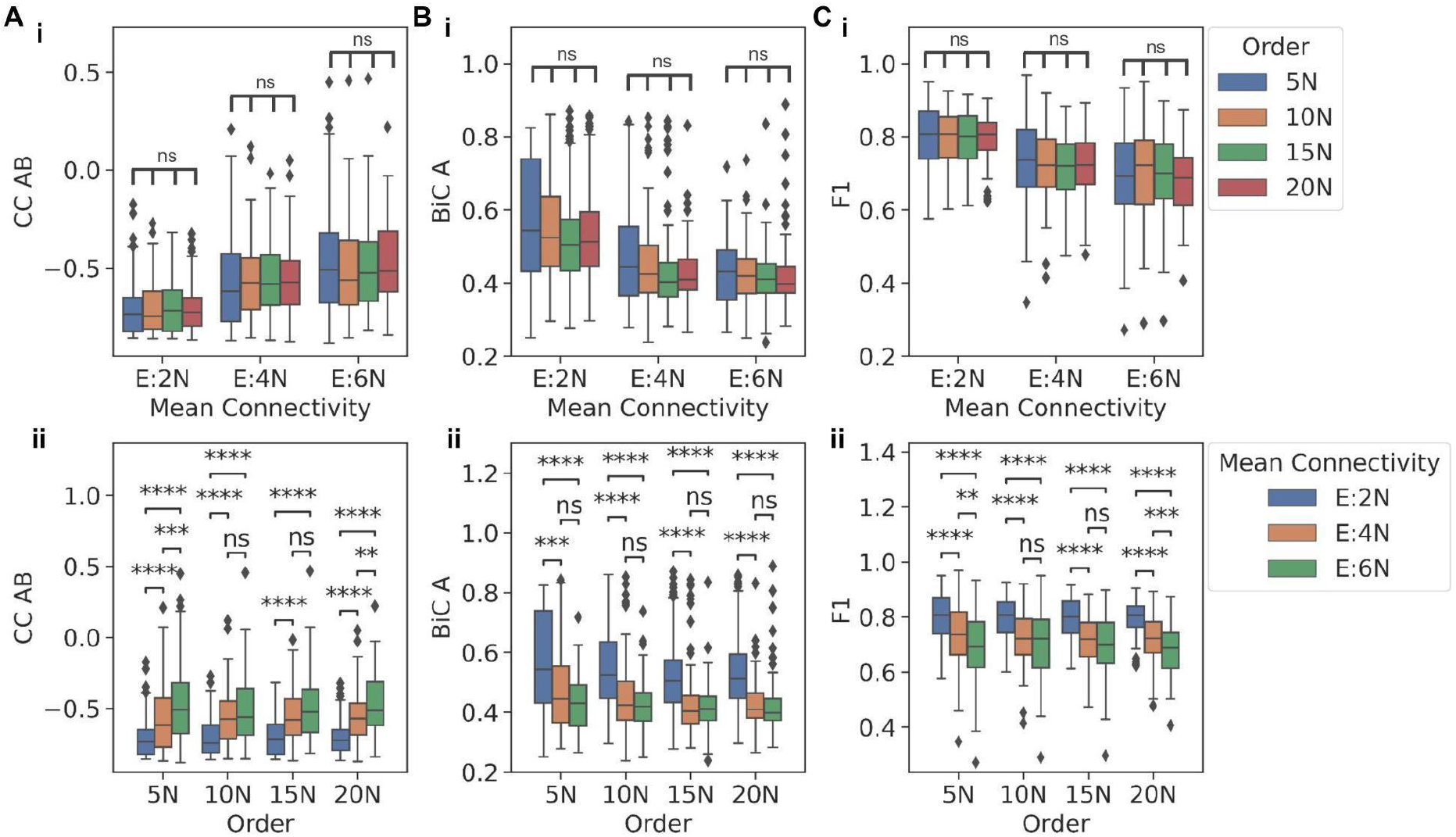
Functional traits of TS embedded in large networks. **A)** Comparison between the distributions of CC AB for TS embedded in: i) networks of same mean connectivity but having different orders and (i) networks of same order having different mean connectivity. **B)** Comparison between the distributions of BiC A for TS embedded in: i) networks of same mean connectivity but having different orders and ii) networks of same order having different mean connectivity. **C)** Comparison between the distributions of F1 for TS embedded in: i) networks of same mean connectivity but having different orders and ii) networks of same order having different mean connectivity. p-values of pairwise Mann-Whitney U tests are denoted by: ns - p <= 1, * - 0.01< p <= 0.05, ** - 0.001 < p <= 0.01, *** - 0.0001 < p <= 0.001, **** - p <= 0.0001

For some cases, the decrease observed in metrics between E:4N and E:6N mean connectivity values was not significantly different, potentially because the TS dynamics was compromised enough in the E:4N case but not in the E:2N case. Put together, all the three metrics considered here to capture the behavior of a TS - CC (how strongly are the two nodes in a TS anti-correlated), BiC (how clearly the high and low levels of a node are segregated), and F1 (how strong is the dominance of mutually exclusivity of the two nodes) - tend to show trends indicating a weakening of the dynamical behavior of a TS, as it is embedded in increasingly denser large networks.

### Local density around a toggle switch impacts its dynamic behavior

Mean connectivity of a network is the ratio of the total number of edges to the number of nodes in the network, i.e. a measure of global network density. Thus, assuming that the network, on average, is equally sparse or dense, with an increase in mean connectivity of the network, the average in-degree of the nodes of a TS embedded in the network also increases. To ascertain whether this increase in the in-degree (in A and in B) for the TS nodes (as a consequence of the increased mean connectivity of the network) contributed to divergence from stand-alone TS dynamics, we analyzed the variation in the three metrics (CC, BiC, and F1) with a change in the in-degrees of the TS nodes. We observed that as the in-degree of both nodes of a TS increased, the mean CC AB values decreased in magnitude, i.e. the TS nodes A and B were not as strongly negatively correlated with one another as in a stand-alone case (**Fig 4A, i**). For an in-degree of one for both the nodes in a TS, i.e. the case when the two nodes only had outgoing edges apart from their mutual inhibitions, we noticed a mean CC AB value of -0.83 (**Fig 4A, i**), the same as that of an isolated TS motif (**Fig 1B**). Furthermore, the magnitude of CC AB showed the fastest decline when both the nodes had equally increasing in-degrees (along the diagonal of the heatmap shown in **Fig 4A, i**). Similarly, F1 decreased steadily with in-degrees increasing equally for the two nodes (**Fig 4A, ii**). On the other hand, the BiC of a given node in TS changed only with the in-degree for that node, and not with the in-degree for the other node, or with overall in-degree of a TS (**Fig S2A: i, ii**). Additionally, the mean BiC values for nodes with in-degree more than two were lower than the typical cut-offs considered for bimodality (∼0.55), indicating the compromised canonical bimodal distributions observed in the nodes of an isolated TS.

**Figure 4:**
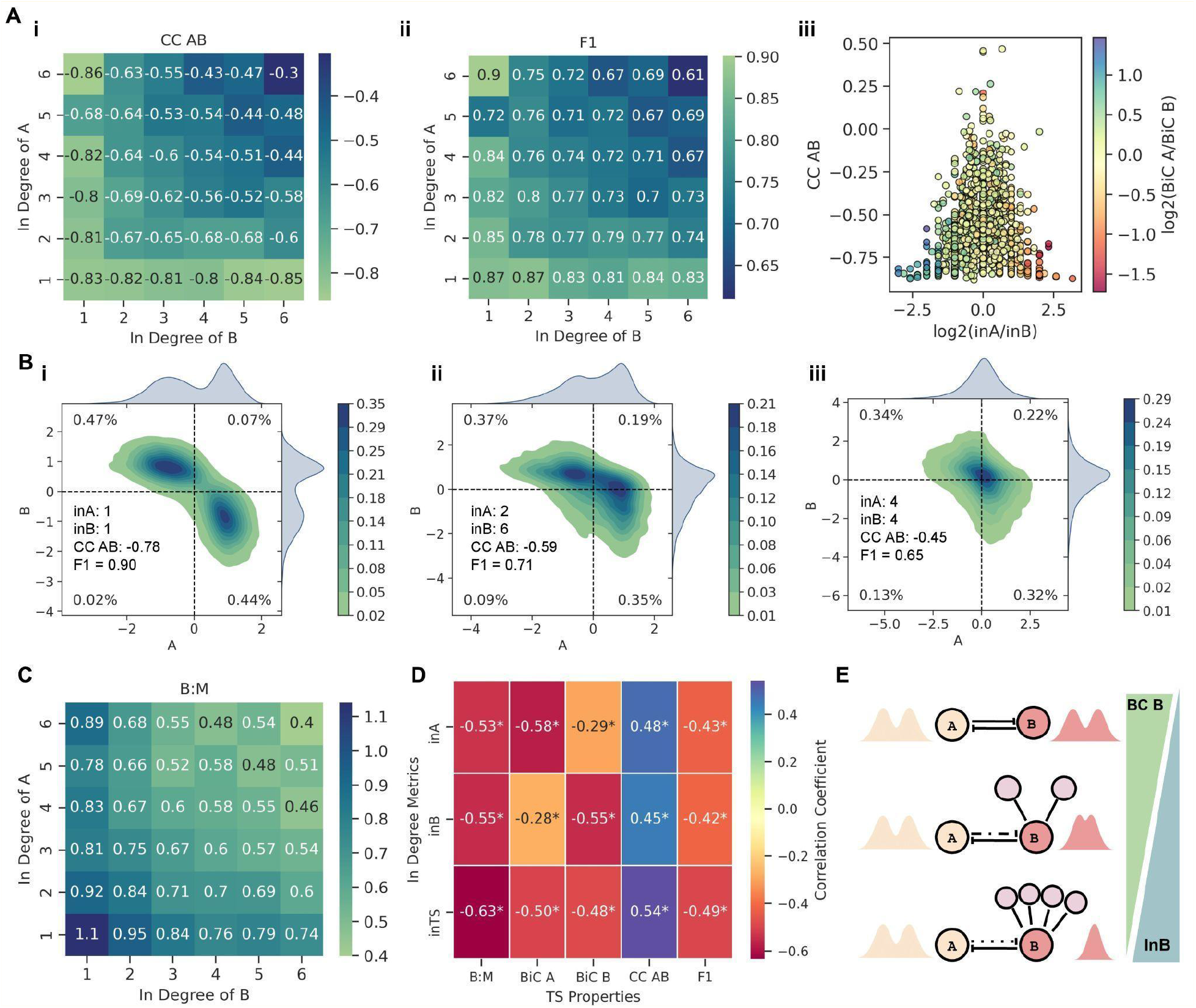
Influence of in-degree of TS on its functional traits. **A)** Heatmaps of values of i) CC AB and ii) F1 for varying in-degree of the two nodes in a TS. iii) Scatter plot of log_2_ (in A/in B) values against their CC AB values, with points colored according to their log_2_ (BiC A/BiC B) values. **B)** Bivariate plots of steady state values of TS nodes with varying in-degree ratios. **C)** Heatmap of the ratio of the fraction of bistable parameter sets which show ‘single-positive’ (01 or 10) steady states to the fraction of monostable parameter sets showing 01 or 10 steady states (B:M), with varying in-degree combinations for the two nodes of a TS. **D)** Heatmap of the pairwise correlation coefficients of in-degree metrics, in-degree of A (in A), in-degree of B (in B) and in-degree of TS (in TS) against various TS properties. *: p-value < 0.05 according to Spearman correlation test. **E)** Summary of the effect of local density on the dynamics of an embedded TS motif.

Further, we quantified the impact of asymmetry in terms of incoming edges on a TS by considering the impact of the ratio of in-degree of A to that of B (log_2_ (in A/in B)) on CC AB and relative BiC values simultaneously. We noticed that the higher the asymmetry in terms of in-degree (log_2_ (in A /in B) > 1 or log_2_ (in A/in B) < -1), the stronger the negative correlation between the two TS nodes (CC AB < -0.5) (**Fig 4A, iii**). Importantly, as the magnitude of log_2_ (in A/in B) increased, the range of CC AB values (initially even spanning positive values; above the red dotted horizontal line), narrowed to highly negative values close to those observed in isolated TS nodes (−0.83) (**Fig 4A, iii**). Similarly, the value of F1 approached closer to that observed in an isolated TS, as the magnitude of (log_2_ (in A/in B)) increased (**Fig S2B, i**). Also, we noted that the higher the in-degree of a node, the more likely it becomes for that node to lose its bimodality seen in a stand-alone TS (log_2_ (BiC A/BiC B) < -1 for log_2_ (in A /in B) > 1 and log_2_ (BiC A/BiC B) > 1 for log_2_ (in A/in B) < -1) **(Fig 4A, iii)**. Together, this analysis suggest that while F1 (fraction of single positive states) and CC (AB) (correlation coefficient) depend on the in-degree of a TS, the BiC of individual nodes depend on the in-degree of respective node.

To substantiate this trend further, we investigated representative cases of varied in-degrees of A and B. When the TS has only outgoing connections and there is no asymmetry between the in-degree of A and B (in A= in B= 1; the only incoming links on A and B are from each other), the bivariate plot of A and B is very similar to that of an isolated TS (compare Fig 1B with **Fig 4B, i**). But, upon asymmetry in the in-degrees of nodes in a TS (in A=2, in B=6, in TS = 2 + 6 = 8), the node with higher in-degree (B) starts to lose its switch-like behavior and shows a more unimodal distribution of its steady state values (**Fig4B, ii**). However, the strongly negative correlation between the TS nodes and concomitantly the fraction of “single-positive” states does not decrease as sharply compared to those for the stand-alone case (compare F1 and CC (AB) in **Fig 4B, ii** with those in **Fig 4B, i**). To deconvolute the impact of higher in-degree of TS vs. asymmetry in the in-degree of both the nodes, we considered a case with the same in-degree for both nodes, without changing the net in-degree for a TS (in A= 4, in B= 4, in TS = 4 + 4 =8). Here, the switch-like behavior of both nodes is largely lost and they show a unimodal distribution of their respective steady state values (**Fig4B, iii**). However, CC (AB) and F1 are comparable to the case of asymmetric in-degrees (compare CC (AB) and F1 values in **Fig 4B, ii** with those in **Fig 4B, iii)**, further supporting that while the higher the in-degree of a toggle switch, the weaker the negative correlation between nodes and the smaller the fraction of “single-positive” states, the bimodality patterns for each node depend on in-degree of that individual node and not on in-degree for TS.

After looking at these representative trends showcasing an increasing in-degree of a node leading to loss of bimodality in steady state distributions of the corresponding node, we investigated how generic these trends were for embedded TS motifs. We hypothesized that with an increasing in-degree of TS, the frequency of co-occurring ‘single positive’ states (01 and 10 states in a bistable setting) should decrease, with a concomitant increase in having either of these two states, i.e. 01 or 10 states in a monostable setup. This feature can be quantified by the ratio of the fraction of bistable parameter sets showing 01 and 10 steady states to the fraction of monostable parameter sets showing 01 or 10 steady states (B:M). When the TS has only outgoing connections to the network (in A= in B= 1; the only incoming links on A and B are from each other), the B:M ratio is greater than 1 (**Fig 4C**). But, as the in-degree for the TS increases, B:M decreases to values below 1, with the sharpest decline when both nodes have equal in-degrees (along the diagonal of the heatmap in **Fig4C**). These results show that as the in-degree of TS increases, the canonical bistable behavior (co-existing ‘single positive’ states) starts to decrease, and simultaneously, the fraction of monostable ‘singe-positive’ steady states increases, implying a loss of bistable traits of the TS motif.

Finally, across larger networks of varying sizes and mean connectivity values, we interrogated how in-degrees for individual nodes as well as for a TS (In A, In B and In TS) correlates with various metrics – F1, CC AB, BiC A, BiC B and B:M ratio. Net in-degree of the TS (in TS) was found to best explain the decline in the magnitude of CC AB, F1 and the B:M ratio (**Fig 4D**). The bimodality coefficients (BiC A and BiC B), on the other hand, were more influenced by the in-degree of their respective nodes and not inTS (**Fig4D**). Therefore, it is the local density on the TS motif (given by in TS) that drives the divergence from TS-like behavior rather than the properties of the whole network in which a TS is embedded.

### In a toggle triad, the fraction of single-positive states capture its functional resilience

After investigating the patterns seen in a TS embedded in large networks, we focused our attempt to understand the functional resilience of the TT motif. We embedded it into the previously described 12 types of large random networks. Similar to observations in TS, the distributions of pairwise correlation coefficients – CC AB, CC BC and CC AC – did not show any significant consistent variation when they were grouped by mean connectivity and compared across the different network orders (**Fig 5A, i; S3A, i; S3B, i**). Intriguingly, unlike the observations in TS, we did not observe any significant differences when the CC values between the TT nodes were grouped by their network orders and compared across the three mean connectivity either (**Fig 5A, ii; S3A, ii; S3B ii**), despite a visible increase in the range of values. Thus, we investigated how the maximum of the three pairwise correlation values (MaxCC) between the TT nodes – CC AB, CC BC and CC AC – varied as a function of network order and/or mean connectivity. We observed that when grouped by order, the higher the mean connectivity, the higher the average MaxCC value; however, no such trend was seen when grouped by mean connectivity (**Fig 5B, i-ii**). Reminiscent of observations in TS, the fraction of “single-positive” (010, 100, and 001) steady states (F1) also decreased overall when comparisons were made across their mean connectivity (**Fig 5C, i**), but not across their network orders (**Fig 5C, ii**). Consistently, the fraction of “double-positive” (011, 110, 101) states (F2), and the fraction of “all-positive” or “all-negative” (111, 000) states (F3) increased across different mean connectivity values when grouped by network orders (**Fig 5D, i-ii**), but not when grouped by mean connectivity and compared across network orders. (**Fig S4A, i-ii**). The ratio of the fraction of “single-high” to that of “double-high” states (F1/F2) also showed the same trend, asymptotically reaching the value of one (**Fig 5D, iii; S4, iii**). These observations help understand the trend seen for MaxCC tending towards positive values with increasing mean connectivity. With decreasing frequency of “single-positive” states, the negative pairwise correlation between the TT nodes starts to weaken, and in some cases, one or more of the correlation coefficient values can be positive, suggesting a decay of stand-alone TT dynamics.

**Figure 5:**
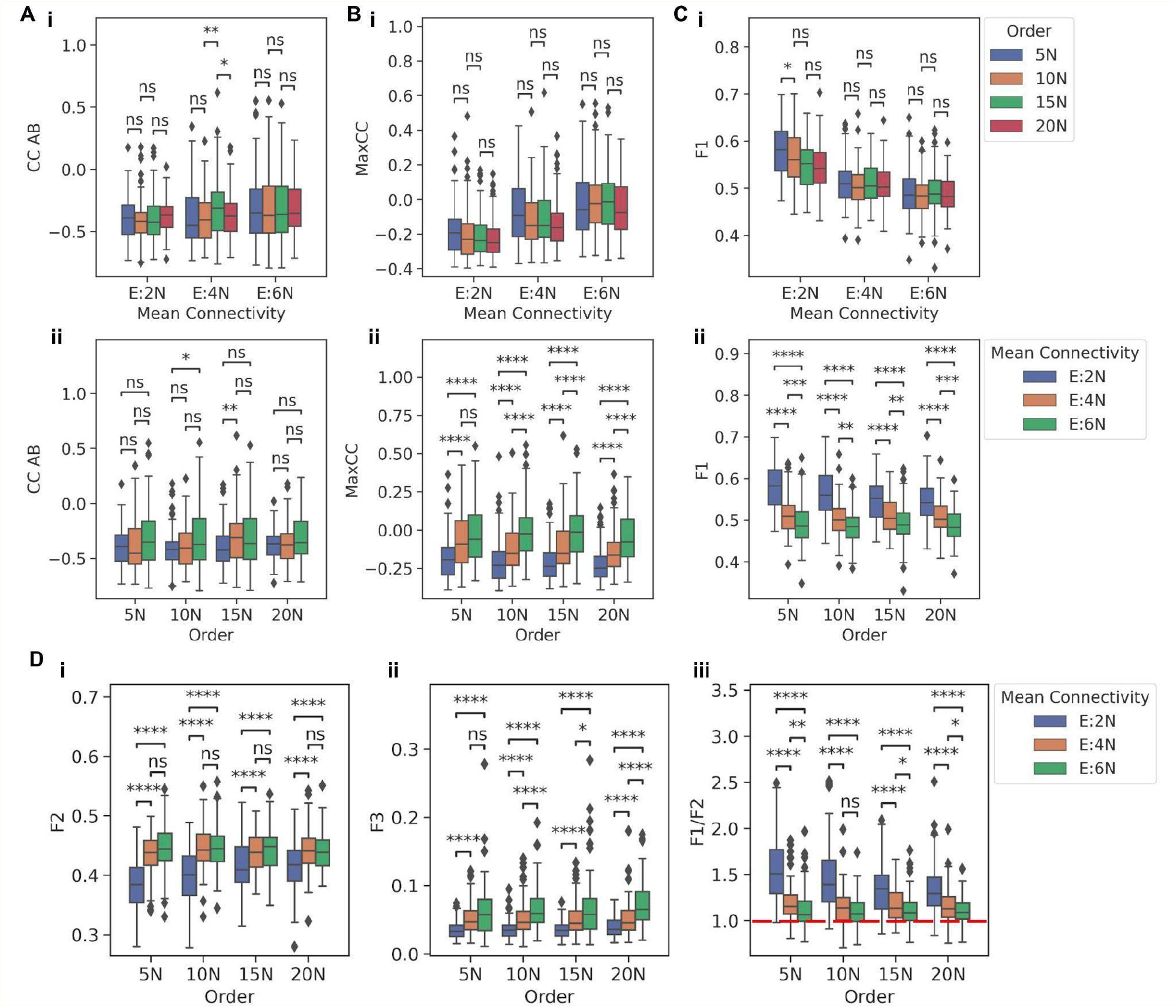
Functional traits of TT embedded in larger networks. **A)** Comparison between distributions of CC AB for TT embedded in i) networks of same mean connectivity but having different orders and ii) networks of same order having different mean connectivity. **B)** Comparison between the distributions of MaxCC for TT embedded in i) networks of same mean connectivity but having different orders and ii) networks of same order having different mean connectivity. **C)** Comparison between the distributions of the F1, for TT embedded in (i) networks of same mean connectivity but having different orders and (ii) networks of same order having different mean connectivity. **D)** Comparison of the distributions of i) F2 ii) F3 iii) F1/F2 for TT embedded in networks of the same order having different mean connectivity. p-values of pairwise Mann-Whitney U tests are denoted by: ns - p <= 1, * - 0.01< p <= 0.05, ** - 0.001 < p <= 0.01, *** - 0.0001 < p <= 0.001, **** - p <= 0.0001

We next investigated how the in-degree of the embedded TT motif affected its behavior. MaxCC values correlated positively (ρ = 0.41, p < 0.05) with the in-degree of TT, indicating that the higher the in-degree of TT, the stronger the decay of TT dynamics (**Fig 6A, i**). Consistently, F1/F2 values decreased as the in-degree of TT increased (ρ = -0.51, p < 0.05), tending towards a value of 1 for high in-degrees of TT (**Fig 6A, ii**), driven by decrease in F1 and increase in F2 and F3 (**Fig S5A**). Moreover, Max CC correlated negatively with F1 (ρ = -0.48, p < 0.05) and F1/F2 (ρ = -0.45, p < 0.05) but positively with F2 (ρ = 0.37, p < 0.05) and F3 (ρ = 0.35, p < 0.05) (**Fig 6A, iii; S5B**). Therefore, with an increasing in-degree of a TT, the fraction of “single-positive” states diminish as they are replaced by “double-positive” (and, to some extent, by “all-positive” or “all-negative”) states. Subsequently, this change in frequencies of different states can weaken the canonical mutual inhibition among nodes in a stand-alone TT, driving one or more of pairwise correlation values to be positive, thus validating our choice of Max CC as a metric to assess the decay of TT dynamics.

**Figure 6:**
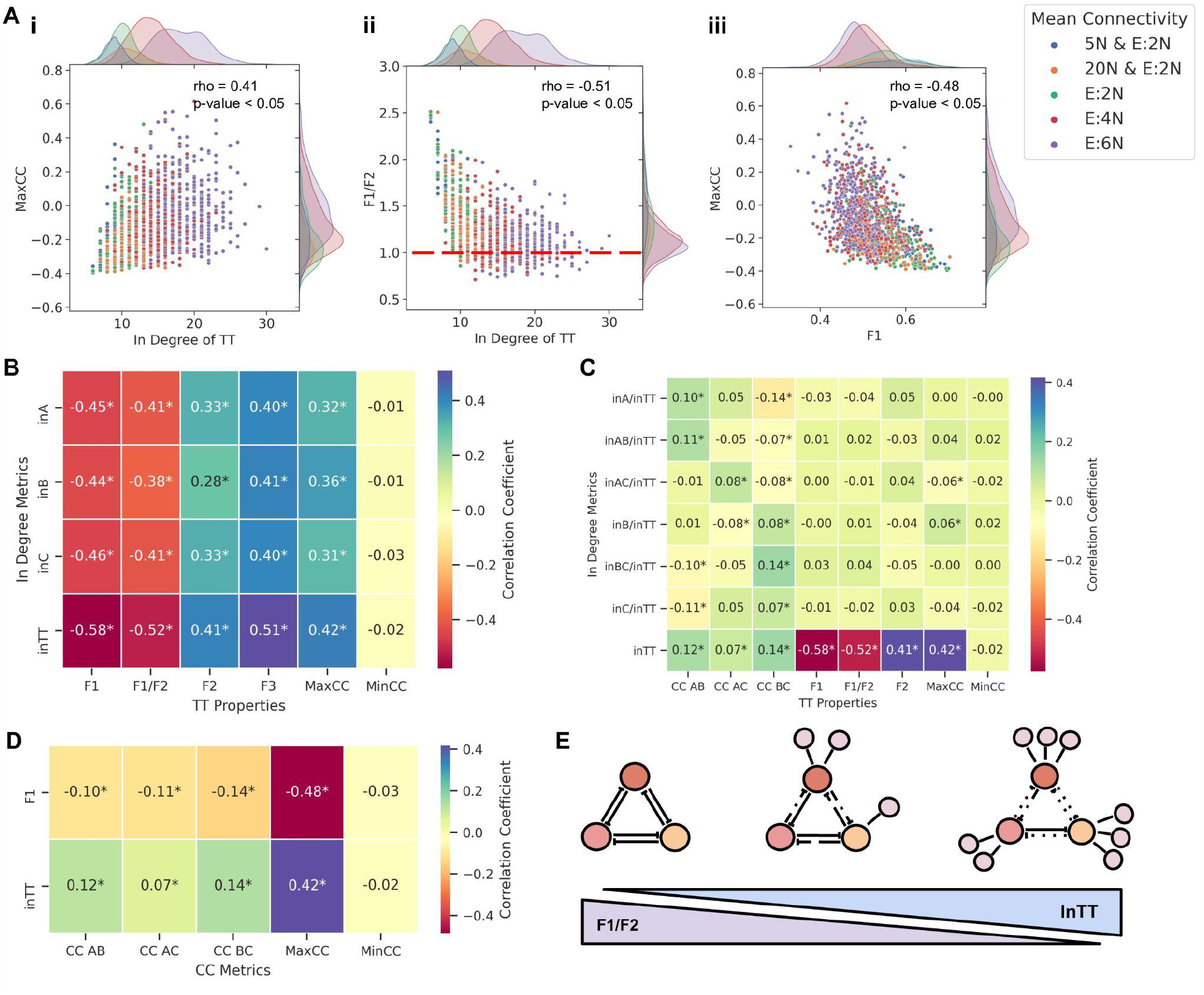
Influence of the in-degree of TT on its dynamics. **A)** i) Plot showing the dependence of change in distribution of the MaxCC values with changing in-degree of a TT motif. ii) Plots showing the dependence of change in the distribution of F1/F2 values with changing in-degree of a TT motif iii) Plots showing the dependence of change in the distribution of MaxCC with changing F1 values. Each dot is colored according to its respective network mean connectivity. Spearman correlation coefficients (rho) and p-values are given in upper right corner of each plot. **B)** Heatmap of the pairwise correlation coefficients of in-degree metrics against various TS properties. **C)** Heatmap of pairwise correlation coefficients of normalized in-degree metrics against various TS properties. In panels B and C. **D)** Heatmap of the pairwise correlation coefficients of inTT and F1 against various CC metrics. * in the heatmap cells signify p-value < 0.05 according to Spearman correlation test. **E)** Schematic summary of the effect of increase in-degree on the dynamics of a TT.

After characterizing the effect of net in-degree of TT on its behavior, we investigated how the three pairwise correlation between the TT nodes (CC AB, CC BC, and CC AC) varied with varying in-degrees for the nodes. Unlike the observations for embedded TS where the CC AB decreased with network mean connectivity and in-degree for motif nodes (**Fig 4A, i; S6**), the pairwise CCs did not show any discernible trend in an embedded TT, therefore, we excluded them from any further analysis. On the other hand, F1, F2 and F3 changed significantly as the in-degrees of any two nodes in the TT increased (F1 decreased while F2 and F3 increased); at high in-degrees for any two nodes, F2 is approximately equal to F1 and 6-7 times the corresponding F3 values (**Fig S7**). Next, we quantified changes in these metrics brought about by simultaneously varying the in-degree of the third node as well. An increase in in C, while maintaining the values of in A and in B, led to a lower F1, higher F2 and F3 values, as expected (compare corresponding cells in **Fig S8B** with **Fig S8A**), thereby showcasing the impact of increasing net in-degree on the breakdown of stand-alone TT dynamics.

Because pairwise correlation coefficients between TT nodes were unable to gauge the changes in TT dynamics, we performed multiple linear regression (MLR) on steady state values of the TT nodes to understand the changes in inter-node dependence brought about by embedding of the TT. MLR was performed by taking steady state values of A and B as independent variables and steady states of C as the dependent variable. The distributions of coefficients of A and B (ACoeff and BCoeff, respectively, i.e. the slopes of the regression plane) and the model’s scores (R^2^ values) were then compared across network orders by grouping them according to common mean connectivity and *vice versa*. As the mean connectivity of the network increased, the magnitude of the mean values of ACoeff and BCoeff decreased, indicating that the repressing effect of the two nodes on C decreased as the network density increased (**Fig S9A-B, i-ii)**. Also, the scores of MLR models declined as the mean connectivity of the network increased, pointing towards a decline in the predictive power of the MLR model as the TT dynamics deviated from their canonical behavior with the increasing network density (**Fig S9A iii; S9B, iii**). Further, the higher the in-degree of TT, the less the model’s score and the magnitudes of ACoeff and BCoeff values (**Fig S9C**), reiterating the disruptive impact of in-degree of TT on its canonical behavior. Intriguingly, for E:6N, we noticed some ACoeff and BCoeff values to be positive (**Fig S9C**), reminiscent of MaxCC values also becoming greater than zero for high mean connectivity (**Fig 6A, iii**).

Next, we performed a meta-analysis across network orders and mean connectivity in terms of correlation of various metrics with in-degree of individual nodes as well as net in-degree of TT. With all metrics considered here (F1, F2, F3, Max CC, F1/F2), In TT showed a stronger correlation (ρ > 0 for F2, F3; ρ < 0 for F1, F2/F2, MaxCC) than the in-degree of any of the individual nodes (in A, in B, in C), indicating that net in-degree was the best predictor of embedded TT dynamics. (**Fig 6B**), and reinforcing the trends seen in embedded TS (**Fig 4D**). This trend was consistent even when the individual in-degree or combined in-degree of two nodes, normalized by net in-degree (in A/in TT, in B/in TT, in C/in TT, in AB/in TT, in BC/in TT, in AC/in TT) was considered (**Fig 6C**). Intriguingly, while in TT correlated strongly with MaxCC (i.e. the pair of nodes whose “mutual exclusion” is the weakest), it did not correlate as strongly with individual pairwise correlation coefficient (CC AB, CC BC, C AC) (**Fig 6C**). Also, while MaxCC correlated strongly with individual in-degree as well as with net in-degree, Min CC (the minimum value among three pairwise correlation coefficients, i.e. the pair of nodes showing strongest mutual inhibition properties) did not show any such association (**Fig 6B, C**). This difference in terms of MaxCC vs. MinCC is consistent with observations that all three individual pairwise correlation coefficients show only a weak correlation with both in-degree of TT and with F1 (**Fig 6D**), thus justifying our choice of using MaxCC as a metric to track the dynamics of embedded TT.

Put together, these results highlight that while the total in-degree of a TT motif is a good indicator of the decay of TT dynamics (decrease in “single-positive” states and simultaneous increase in “double-positive”), it often does not contain precise information on which out of the three possible pairs of nodes in a TT have their mutual repression compromised and thus drive a decay in the stand-alone dynamical behavior of a TT. Despite this limitation, similar to the results seen for TS, the dynamics of embedded TT was determined not by network order, network density, or individual in-degrees of the motif nodes, but by the total in-degree of the motif (In TT) (**Fig 6E**). Moreover, both for the TS and TT motifs, the nature of these in-degrees - being either activating or repressing - did not affect the nature of divergence from canonical behavior (**Fig S10**), highlighting that the motif is sensitive to total number of incoming edges, not their distributions in terms of their sign/effect.

### Effect of self-activation and self-inhibition of nodes on the modularity of motifs

Many “master regulators” often driving cell-fate decisions and forming a TS or TT can self-activate ^2,13^. To evaluate the effects of self-activations and self-inhibitions on the motif nodes, the TS and TT motifs containing self-activation (TS-SA and TT-SA) and self-inhibition edges (TS-SI and TT-SI) on all nodes were embedded into combinations of two network orders (5N and 20N) and two mean connectivity (E:2N and E:6N), resulting in four types of networks. The same pipeline as before was followed to generate the networks, with n=100 for each type of network (i.e. 400 networks per motif). The motifs TS-SA, TS-SI, TT-SA and TT-SI were then embedded into these networks and simulated with RACIPE, and corresponding metrics were compared. First, we compared the TS, TS-SA and TS-SI embedded in networks having the same mean connectivity. For all the three metrics (CC AB, F1 and BiC), TS, TS-SA and TS-SI showed significantly different distributions (**Fig 7A, i-iii**). While including self-activation moderately strengthened the correlation between A and B, self-inhibition significantly weakened it (**Fig 7A, i**). Similarly, while including self-activation led to slightly increased F1 and BiC values, conversely, including self-inhibition on TS nodes reduced the F1 and BiC values (**Fig 7A, ii-iii**). These results suggest that while adding self-activation on nodes can preserve dynamical features of a TS embedded in large networks, self-inhibition can accelerate the decay of TS dynamics, possibly offering a reason for observed higher frequency of self-activating ‘master regulators’ rather than the self-inhibiting ones ^25^.

**Figure 7:**
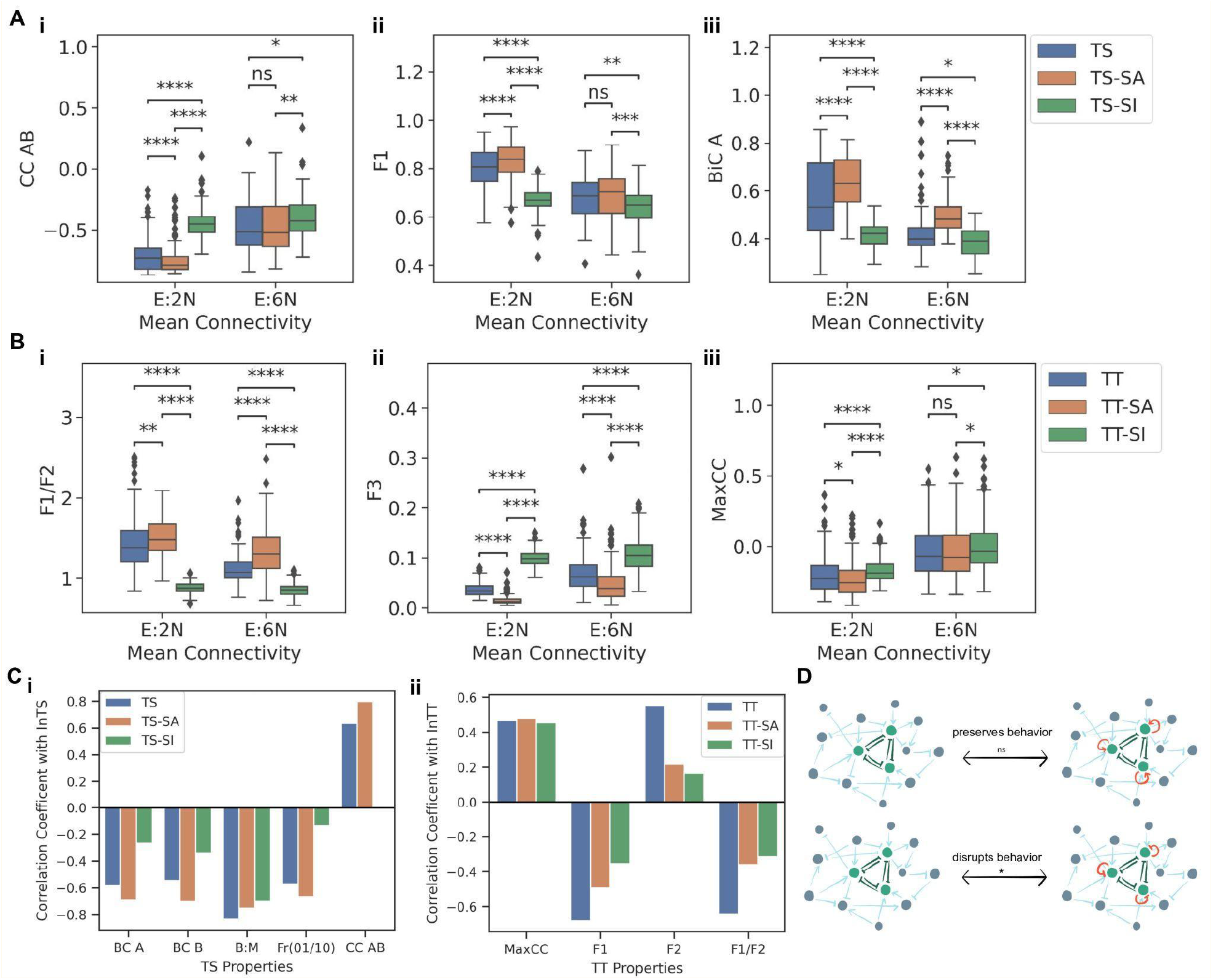
Effect of self-activating and self-inhibiting loops on modularity of TS and TT motifs. **A)** Comparison between the distributions of i) CC AB, ii) fraction of 01 and 10 steady states, iii) BiC A for TS, TS-SA and TS-SI when embedded in networks of same mean connectivity. **B)** Comparison between the distributions of i) F1/F2, ii) MaxCC, iii) F3 for TT, TT-SA and TT-SI when embedded in networks of same mean connectivity. **C)** Comparison of the correlation coefficients of different properties of regular motifs, motifs with self-activation (SA) and self-inhibition (SI) against their respective in-degrees i) For TS ii) For TT. **D)** Schematic showing the effect of self-activating and self-inhibiting edges on motif nodes when embedded in larger networks.

Next, we probed the impact of self-activation and self-inhibition in the case of an embedded TT. Similar to observations in TS, self-inhibition had an opposite and a stronger impact in influencing the metrics as compared to self-activation. Median F1/F2 values noted for TT-SA was higher than those for both TT and TT-SI, irrespective of the mean connectivity (**Fig 7B, i**). Consistently, median F3 values are higher for TT-SI than for TT and TT-SA (**Fig 7B, ii**). Together, these results indicate that while adding self-activation on nodes of a TT can enrich for canonical “single-positive states”, adding self-inhibition can enrich for “all-positive” or “all-negative” states instead. Similarly, for Max CC, TT-SI had higher median values than both TT and TT-SA cases (**Fig 7B, iii**), resulting in faster decay of canonical properties of TT. In other words, the self-inhibiting “master regulators” can exhibit compromised phenotypic decision-making as compared to non-self-inhibiting ones.

Finally, we assessed whether adding self-regulation preserves the correlation of in-degree of motif (TS or TT) with various metrics (BiC A, BiC B, B:M, F1 and CC AB for TS; F1, F2, F1/F2 and MaxCC for TT). We consistently observed that motifs with self-inhibition had lower magnitude of correlation with all these motif properties as compared to motifs with no self-regulation or with self-activation (**Fig 7C, i-ii**), indicating a faster loss and subsequent saturation of motif’s dynamical properties. These trends reaffirm that the addition of self-activating loops preserves the canonical behavior of the motif while adding self-inhibition loops diminishes it remarkably.

## Discussion

Investigating the modularity of biological networks has been an active area of research. Modularity has been loosely defined as corresponding to a highly interconnected set of nodes such that density of connections within that module is significantly higher than that of density of connections of this module with other modules ^26^. Thus, modularity has been mostly studied from a network topology perspective, rather than a functional one. Similarly, the concept of network motifs - recurring sets of regulatory interactions that appear more frequently than expected in a given network - also highlights network sub-structures based on their topology ^27^. While the dynamics of such motifs has received extensive attention ^27^; how insular or intact the dynamics of these network motifs are when embedded in a large network, remains largely underexplored.

Here, we investigated how ‘modular’ the behavior of a TS or TT is when embedded in random large networks of varying sizes and densities. For both these motifs, we observed that an increase in local density around them (i.e., number of incoming edges on TS or TT) was capable of changing their dynamical behavior rather than any global topological properties associated with the large network. Although we witnessed how increasing mean connectivity of the network changed the distributions of the metrics we have used to characterize the dynamical behavior of TS or TT, further analysis revealed that it was the increasing in-degree of the motif (reflected, in part, by mean connectivity as well) that was driving this change in motif behavior. For the TS, it was found that all the three metrics, BiC, CC, and the fraction of ‘single-positive’ (01, 10) states were suitable to gauge its change in dynamics. On the other hand, for a TT motif, CC metrics were not suitable to gauge its behavior, as they did not show extensive variation upon being embedded in large networks. This trend could be because CC, being a pairwise metric, is not optimal to capture the variations in the steady state values of all three nodes. On the other hand, the maximum value of all the 3 pairwise correlation coefficients between the TT nodes, MaxCC, was found to be suitable to gauge the change in dynamics of TT. MaxCC values being correlated positively with double-positive states (F2), was able to capture the enrichment of these states as the in-degree of TT increased. Similar to TS, the fraction of ‘single-positive’ states was also found to be a good metric to observe the variations in the dynamics of an embedded TT. We found that as the in-degree of an embedded TT increased, the single-positive states and double-positive states were almost comparable in frequency. This observation suggests that in the case of CD4+ T-helper cell differentiation - a case study of TT dynamics - the “double positive” cell-states (hybrid Th1/Th2, Th1/Th17, and Th2/ Th17 phenotypes) seen experimentally ^28,29^ could exist due to the TT between GATA3, RORγT and T-bet being driven by various other stimuli that impinge on these nodes via activation or repression. Therefore, besides self-activation ^2,30^, embedding in large networks can be an additional way to enrich such ‘hybrid’ states, as being increasingly reported in various biological systems ^31–33^, both in cases of TS and TT.

The bistable dynamics of a TS motif have been extensively investigated. The nodes of a TS motif show a bimodal distribution ^34–36^ of their steady states dependent upon a balance between the kinetic parameters of the two nodes ^14,37^. An asymmetry in these parameters bringing an imbalance in inhibitory strength can lead to a one-way cause-effect relation between the two TS nodes, with only one node showing bimodality while the other node showing unimodal behavior ^14,20,34,37^. Besides, an increase in the number of downstream interacting elements can also induce competition between produced proteins to bind to the promoter sites of the TS genes, changing the dynamics of motifs and potentially leading to a loss of bimodality ^38^. Here, we show that the relative in-degrees of the nodes can also contribute to this behavior. A skew in the in-degrees led to a divergence from the bistable behavior of a TS, much like the skew in the kinetic parameters. Thus, our analysis uncovers an important design principle of gene regulatory networks (GRNs) that in order to maintain bistable features for a TS, the in-degree, which represents the number of regulators acting at a given point of time, should be minimal. This observation is reminiscent of previous studies demonstrating that in an *E. coli* transcriptional network, no transcription factor had an in-degree or out-degree greater than two, and this feature played a key role in enabling robustness of the network ^39^. Thus, the in-degrees we report here for robustness are in good agreement with those seen in networks for GRNs of various organisms ^39^. Similarly, another recent study associated dynamical robustness of networks with their low in-degree ^40^. Together, these results can help us design optimal strategies to design and integrate synthetic circuits into GRN of a cell rather than developing stand-alone/isolated modules.

Often, various cellular processes are viewed as generally being robust to noise and perturbations, but in diseased states such as cancer, due to a change in network topology, protein production rates or short decay times can potentially perturb these steady states and encourage the progression of disease into states which are more robust and hard to reverse ^37,41–43^. Our results show that in addition to these factors, the local density can also deviate a motif from its canonical behavior and can hence help us develop better algorithms to identify potential drug targets which are more susceptible to alterations in their local neighborhood, to devise treatment strategies to escape these diseased states ^36,44^. The ability to understand the behavior of these multistable states and modulate them in larger networks can help us potentially traverse through the trajectories set by cellular differentiation branching tree by targeting the adjacent incoming edges of the master regulators, in addition to master regulators themselves, to modify the dynamics of cellular decision-making they are involved in, to reprogram cells into the desired cell fate(s) ^4,36,45^.

## Author contributions

MKJ conceived and supervised the research, PH performed the research. All authors contributed to data analysis and writing of the manuscript.

## Supplementary Information Contents

Contains 13 Supplementary Figures.

## Materials and Methods

### Random Network Generation

A total of 12 types of randomized networks were generated for each motif. These network types were combinations of four network orders, with the number of nodes (N) equal to 5, 10, 15, and 20 and three mean connectivities E:2N, E:4N and E:6N with edge to node ratios E:xN, where the number of edges, E, is x times the number of nodes N (Fig S11A). One hundred random networks were generated for each class of network. Thus, 1200 networks were simulated for each motif (TS and TT) to characterize their properties. This analysis was then repeated in triplicates for statistical tests/comparisons.

A custom python3 script was written to generate random networks of a given size and mean connectivity by creating square null matrices (of order N+y, where N is the order of the network and y the number of nodes in the embedded motif) and later populating it with the motif edges, i.e. for a TS embedded in a 5N, E:4N networks, there were total 7 (= 5 + 4) nodes and 22 (= 5*4+ 2) edges. Activating and inhibiting edges were then randomly added depending on the mean connectivity of the network (Fig S11B-C; S12; S13). It was also ensured that no self-activating and self-inhibition edges were formed in any of the network nodes. Generated networks were checked if they were connected using the *isconnected* function of the *networkx* library of python3, and if found to be not connected, they were replaced by a newly generated network. When duplicate networks were found, they were replaced by newly generated networks to avoid any skew in data due to the repetition of network topologies.

### Random Circuit Perturbation (RACIPE)

Random Circuit Perturbation (RACIPE) formalism generates a system of ordinary differential equations (ODEs) for a given network topology and simulates the ODEs by pooling parameters from a randomized predetermined range to identify dynamical properties of a network topology _23_.

For a node T in the network having P_i_ activating and N_i_ inhibiting nodes with incoming edges, the ODE generated by RACIPE to represent the node T will be given by:

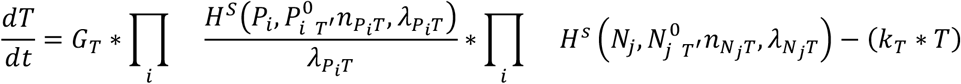

where the terms T, P_i_ and N_i_ are the concentrations of the nodes at time t, n is the Hill coefficient showing the influence of P_i_ or N_i_ on T, λ is the fold change in expression caused by node P_i_ or N_i_ upon acting on node T, P_i_^0^ or N_i_^0^ are the threshold values of Hill function, G_T_ is the production rate, and k_T_ is the degradation rate of the node T.

H^S^ represents the shifted Hill equation and is defined by:

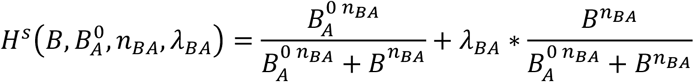

For a particular topology file, RACIPE generates multiple randomized parameter sets and simulates them over multiple initial conditions to identify the steady state levels of the nodes. The parameters are randomized by sampling from their respective pre-defined ranges given below:

Simulations for all the networks were done in triplicates, with 10000 parameter sets per replicate and 100 initial conditions for each parameter set.

**Table 1:** Ranges of randomized parameters in RACIPE

| Parameter | Minimum Value | Maximum Value |
| --- | --- | --- |
| Production Rate (G) | 1 | 100 |
| Degradation Rate (k) | 0.1 | 1 |
| Inhibition Fold Change ($\lambda^-$) | 0.01 | 1 |
| Activating Fold Change ($\lambda^+$) | 1 | 100 |
| Hill's Coefficient (n) | 1 | 6 |
| Threshold | Half-Functional Rule: Sets the threshold depending on the in-degree of the node. |  |

### Normalizing the steady state values

The following formula first normalized the steady state values obtained from RACIPE simulations for each network:

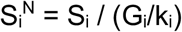

Here, Si^N^ is the normalized steady state value of the i^th^ node; S_i_ is the steady state value of i^th^ node given in RACIPE output; G_i_ is the production rate, and k_i_ is the degradation rate parameter value of the i^th^ node. The normalized values were then converted to z-scores using the *zscore* function of the SciPy library of python3. If a node showed a z-score above zero, it was considered to be showing higher expression and was considered to be in “ON” state represented by 1 and if it showed a z-score below zero, it was inferred to be in “OFF” state represented by 0. This criteria was then used to convert the steady state values into a string of zeros and ones representing the binarized steady state shown for a particular parameter set.

### Sarle’s Bimodality Coefficient (BiC)

Sarle’s bimodality coefficient (BiC) ^24^ was used to identify the nature of the distribution of nodes’ steady state values as either bimodal or not. The formula used to calculate bimodality is given by:

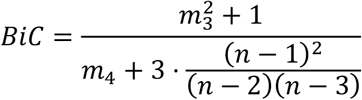

Where m_3_ is the skew of the distribution, m_4_ is the excess kurtosis, and the sample size is denoted by *n*. BiC varies from 0 to 1, with values above 0.55 (i.e. 5/9) representing bimodality in the distribution and those below this threshold indicating a unimodal distribution ^24^. Skew and excess kurtosis were calculated using functions of the SciPy library of python3. The values were then plugged into the above equation to obtain the bimodality coefficients.

### Correlation Coefficient (CC)

Correlation coefficients (CC) was calculated using the function *spearmanr* of the SciPy library of python3.

## Data and Code Availability

The Github repository for RCAIPE-1.0 ^23^ can be found at https://github.com/simonhb1990/RACIPE-1.0.

Custom python3 (python3 version 3.8.10) scripts were also written to analyze the RACIPE output files further and can be found at https://github.com/MoltenEcdysone09/ModularityCodes.

## Supplementary Information

**Figure S1:**
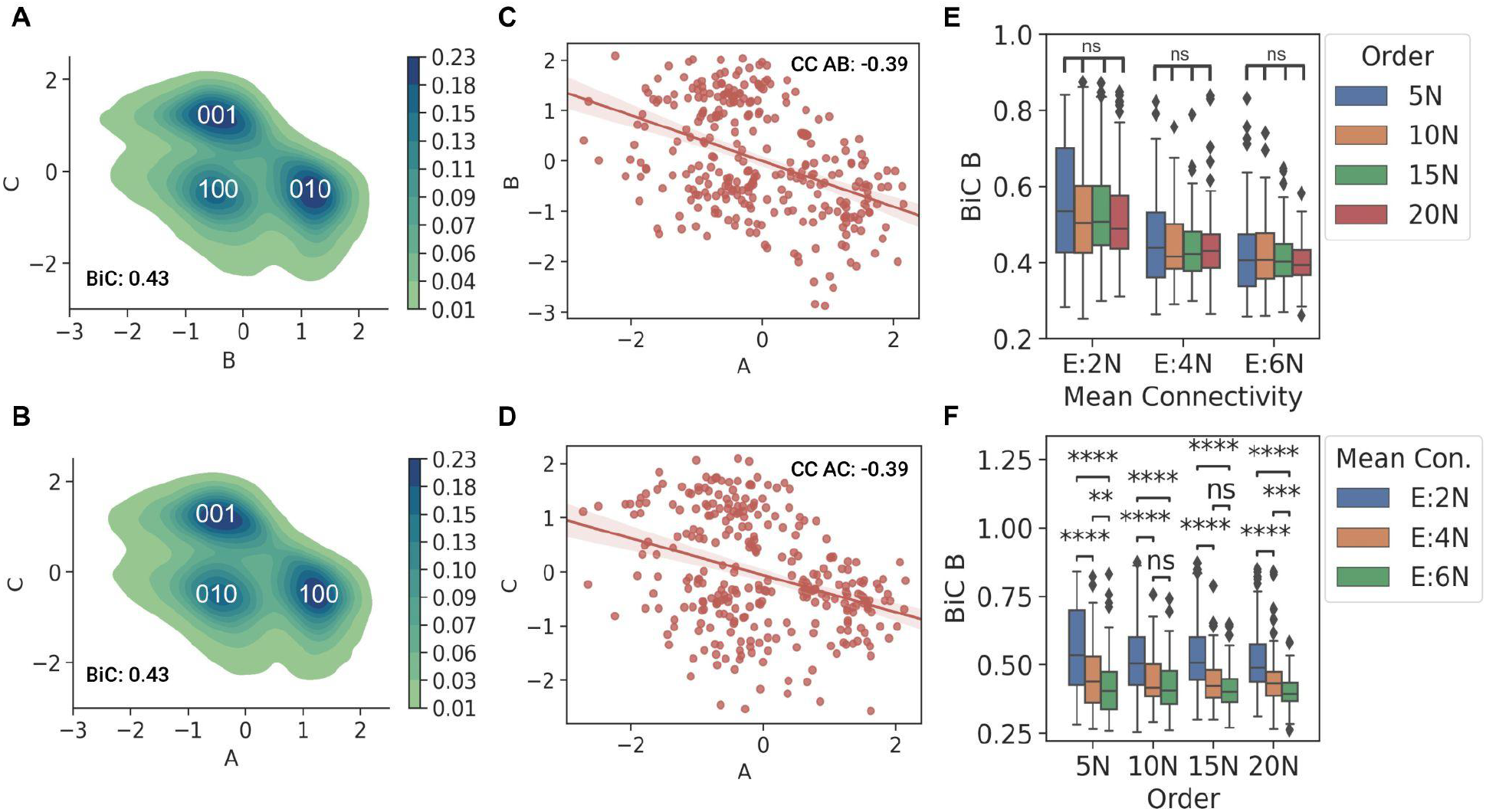
**A)** Probability density plot of steady state values of two TT nodes B and C, the three clusters represent the three single-positive steady states, 001, 010, and 100 shown by the TT motif. **B)** Probability density plot of steady state values of two TT nodes A and C, the three clusters represent the three single-positive steady states, 001, 010, and 100 shown by the TT motif. **C)** Regression plot between steady state values of nodes A and B of a TT. **D)** Regression plot between steady state values of nodes A and C of a TT. **E)** Comparison between the distributions of BiC B for TS embedded in networks of same mean connectivity but having different orders. **F)** Comparison between the distributions of BiC B for TS embedded in networks of the same order but having different mean connectivity. p-values of pairwise Mann-Whitney U tests are denoted by: ns - p > 0.05, * - 0.01< p <= 0.05, ** - 0.001 < p <= 0.01, *** - 0.0001 < p <= 0.001, **** - p <= 0.0001

**Figure S2:**
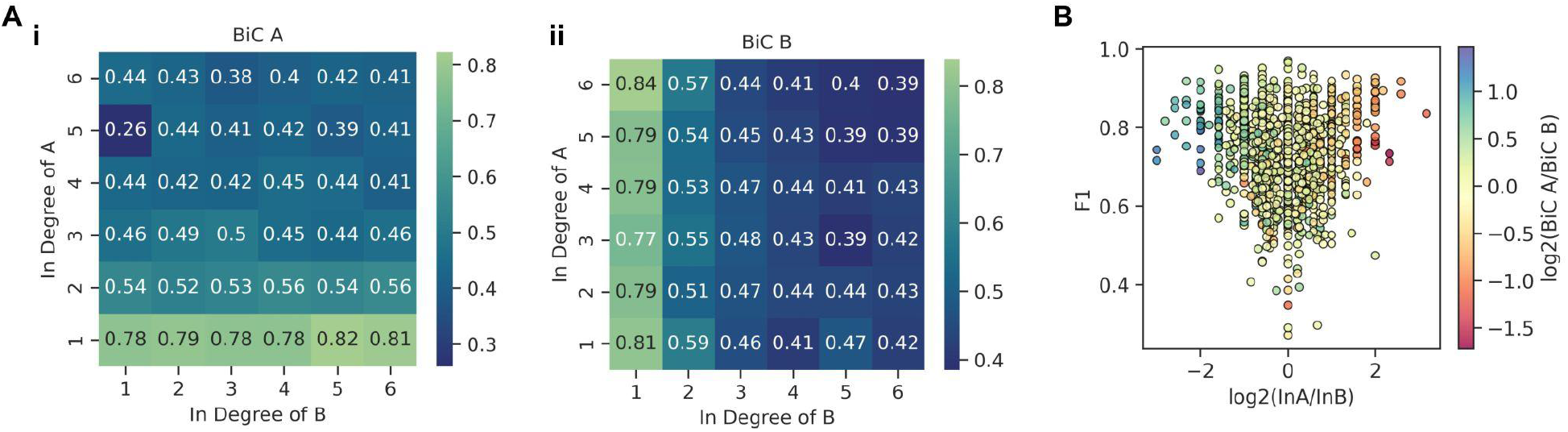
**A)** Heatmap of the variation in (i) BiC A and (ii) BiC B with change in the in-degree of the two nodes of a TS. (B) (i) Scatterplot of the log_2_ (in A/in B) values against F1 values. The points are colored with respect to log_2_ (BiC A/ BiC B) values.

**Figure S3:**
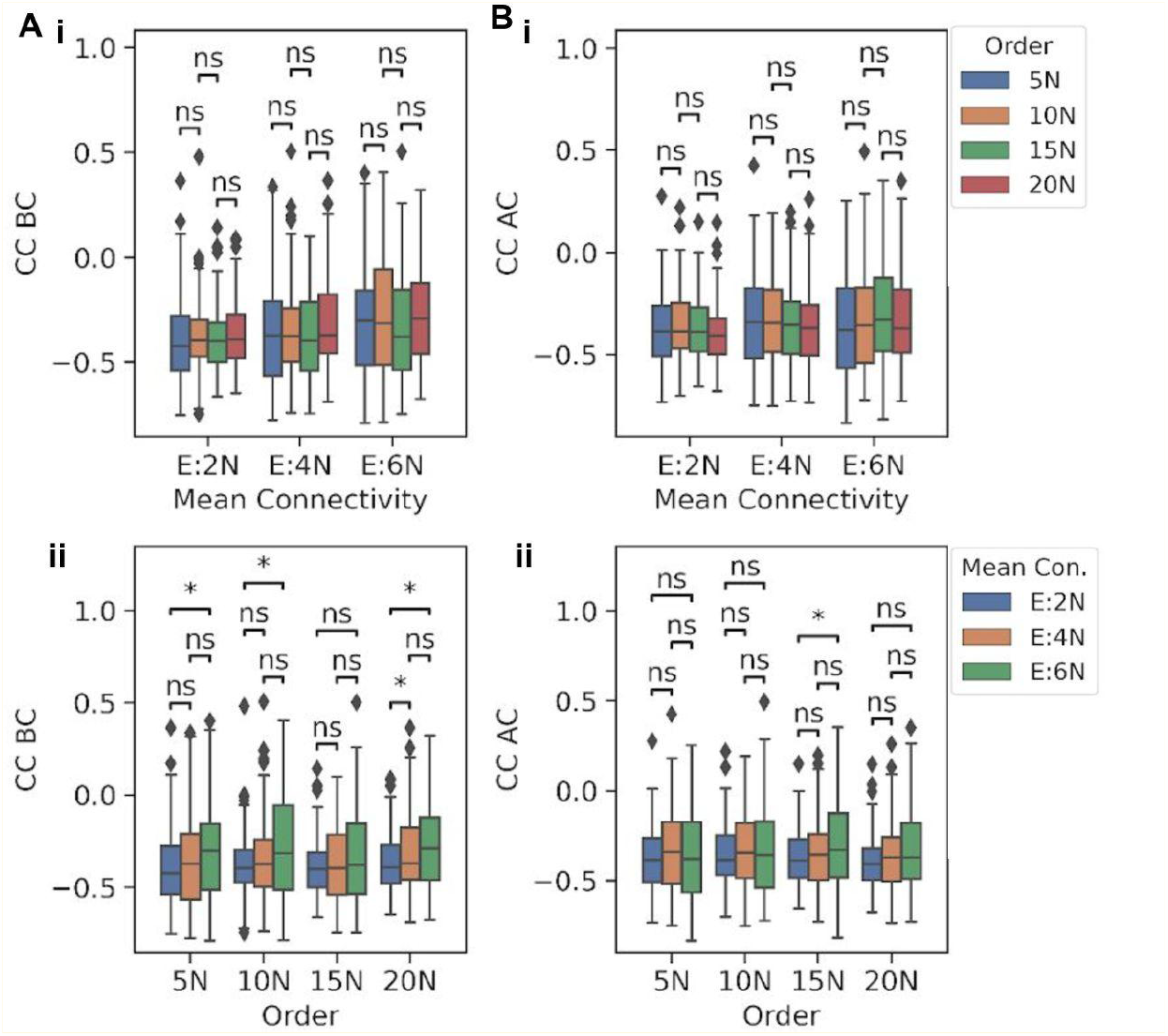
**A)** Comparison between the distributions of a metric for TT embedded in networks of the same mean connectivity but having different orders for: (i) CC BC and (ii) CC AC. **B)** Comparison between the distributions of a metric for TT embedded in networks of the same order but having different mean connectivity for: (i) CC BC and (ii) CC AC. p-values of pairwise Mann-Whitney U tests are denoted by: ns - p > 0.05, * - 0.01< p <= 0.05, ** - 0.001 < p <= 0.01, *** - 0.0001 < p <= 0.001, **** - p <= 0.0001

**Figure S4:**
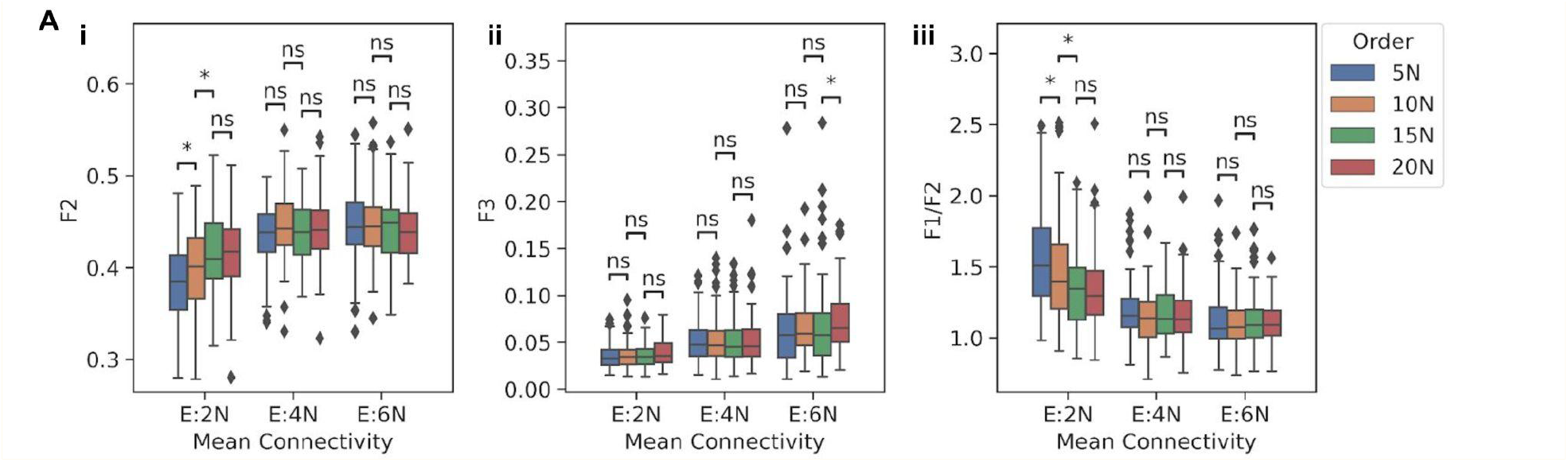
**A)** Comparison between the distributions of a metric for TT embedded in networks of the same mean connectivity but having different orders for: (i) F2, (ii) F3 and (iii) F1/F2.

**Figure S5:**
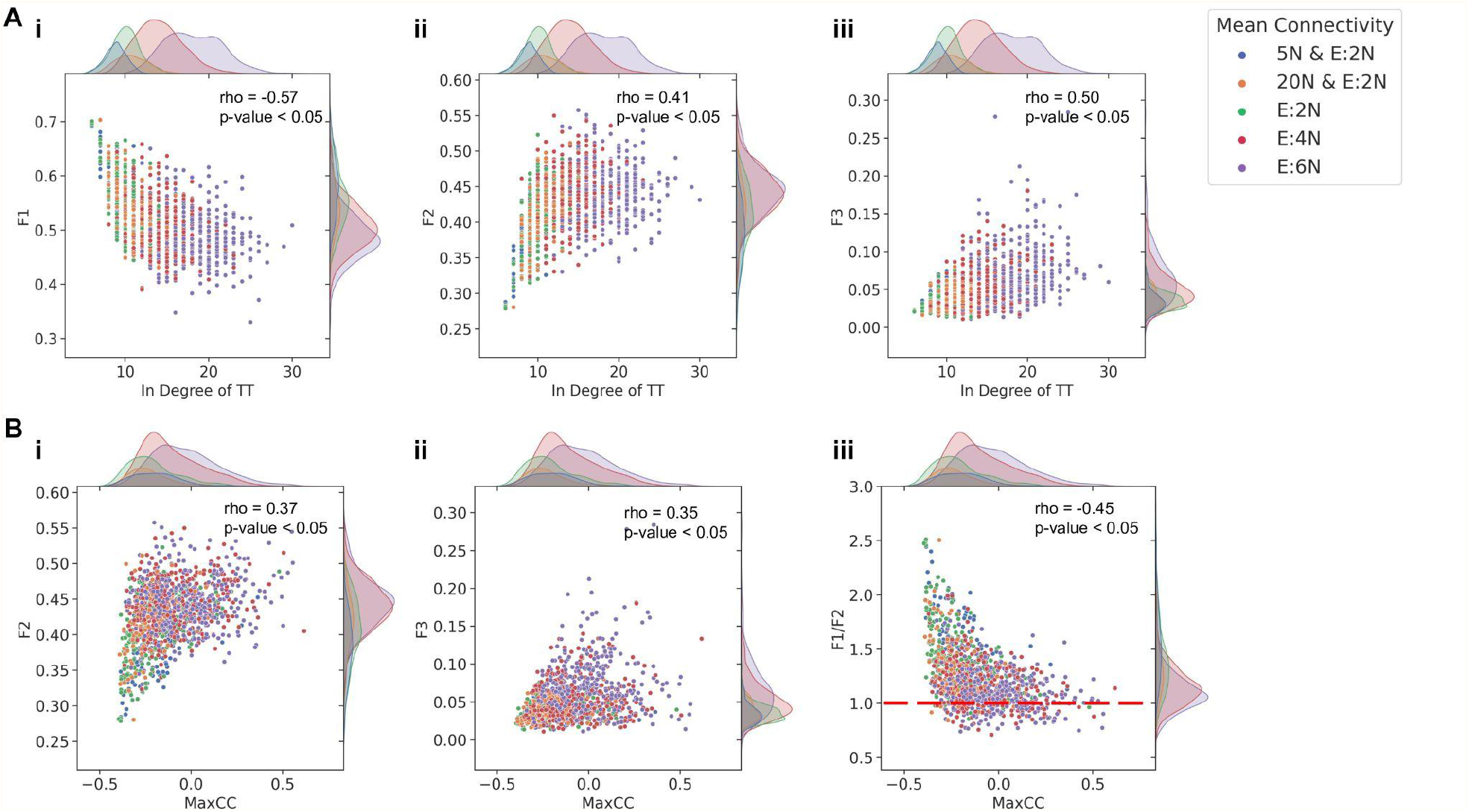
**A)** Plots showing the dependence of change in the distribution of a metric with changing in-degree of a TT, for: (i) F1, (ii) F2, and (iii) F3. **B)** Plots showing the dependence of change in the distribution of a metric with changing MaxCC, for: (i) F2, (ii) F2, and (iii) F1/F2. Each dot is colored according to their respective network mean connectivity values. The spearman correlation coefficients (rho) and p-values are given in the upper right corner of each plot.

**Figure S6:**
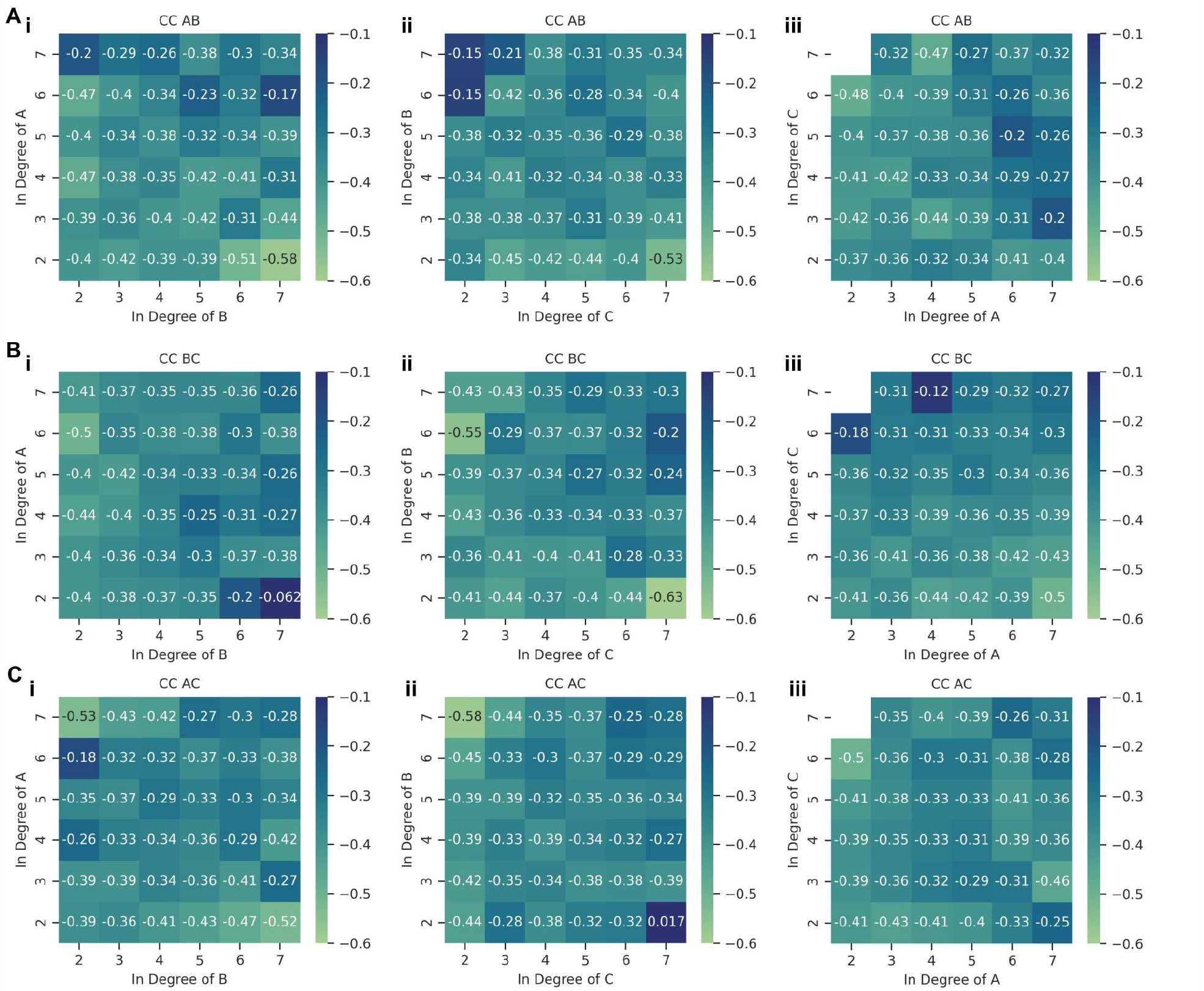
Heatmaps showing the variation of pairwise correlation values against different pairs of in-degrees of TT nodes.

**Figure S7:**
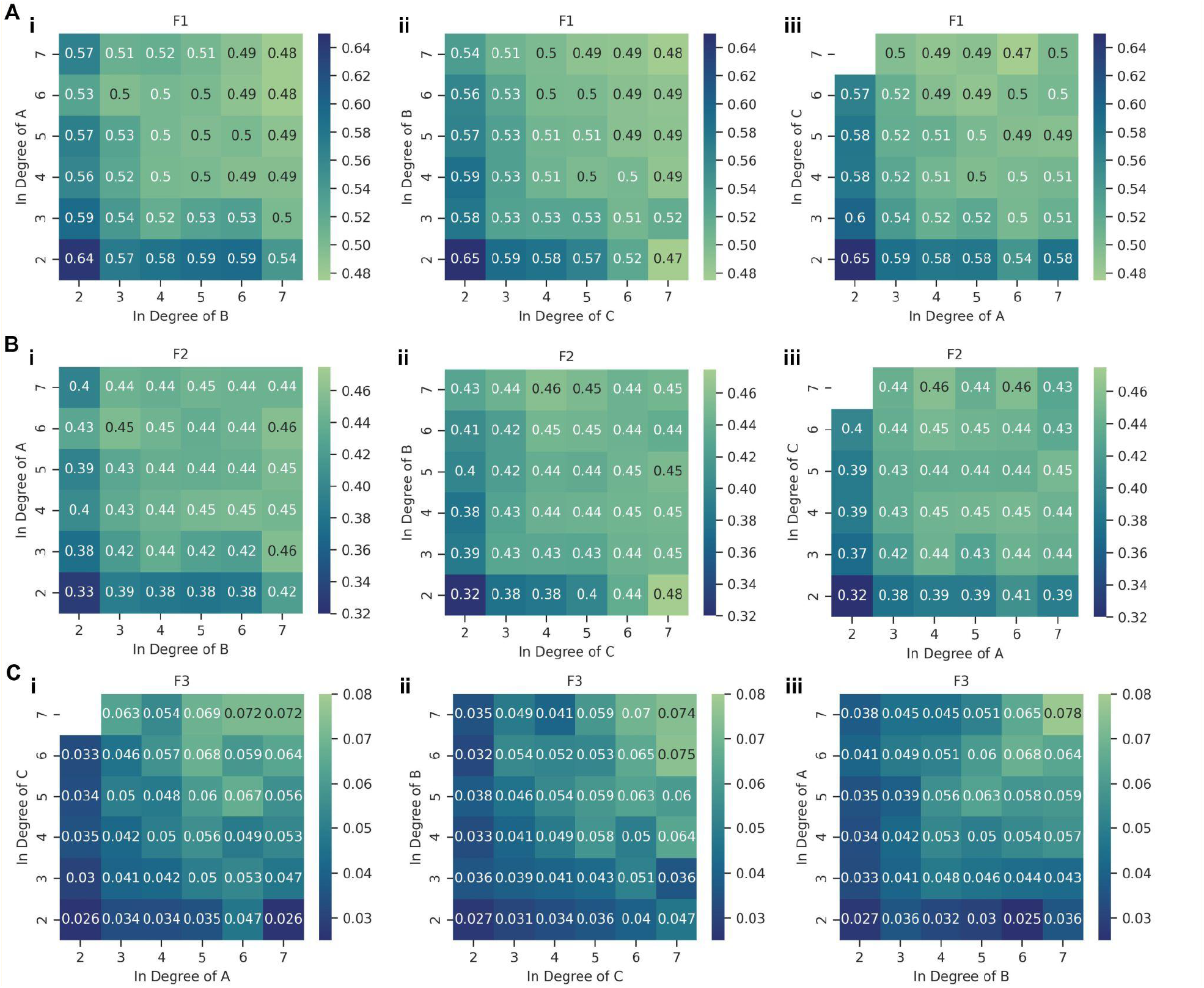
Heatmaps showing the variation of **A)** F1, **B)** F2, and **C)** F3 values against different pairs of in-degrees of TT nodes.

**Figure S8:**
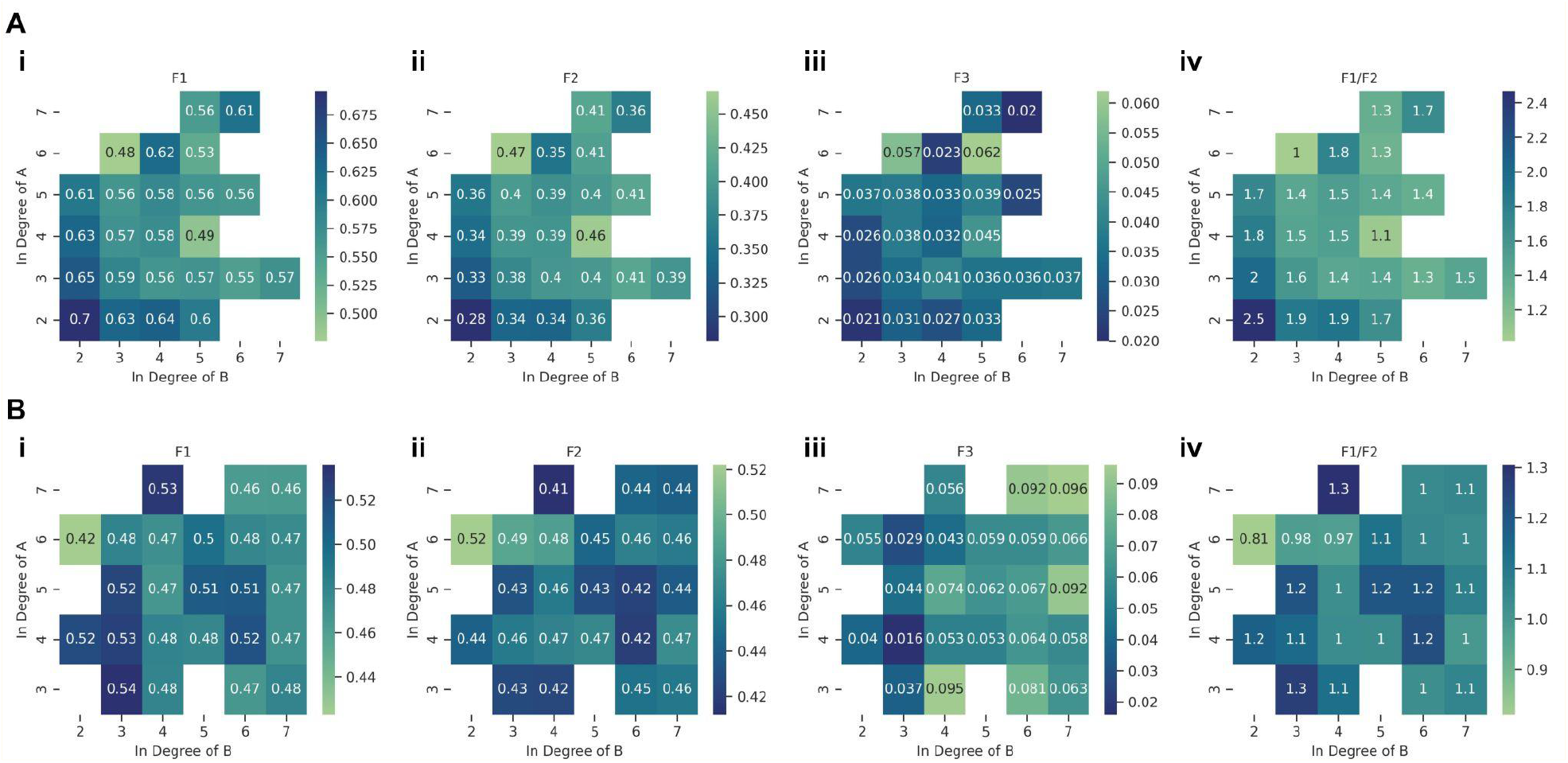
Heatmaps of the variation of F1, F2, F3 and F1/F2 metrics respectively for different combinations of in A and in B for (i) in C = 2 and (ii) in C = 7. Empty cells (shown in white) indicate that no corresponding networks were found in the ensemble of networks we investigated here.

**Figure S9:**
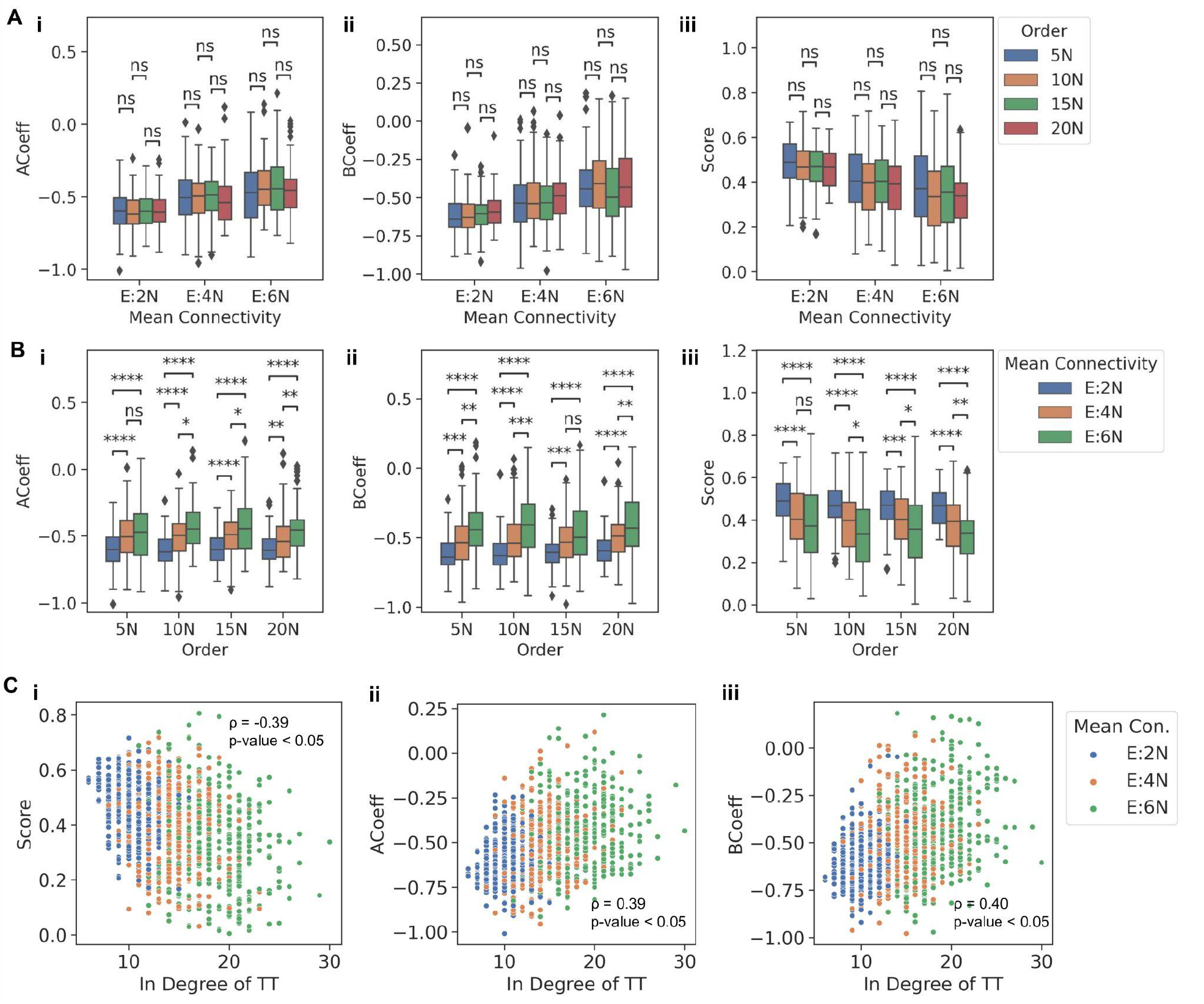
**A)** Comparison between the distributions of a metric for TT embedded in networks of the same mean connectivity but having different orders for: (i) ACoeff, (ii) BCoeff and (iii) Score. **B)** Comparison between the distributions of a metric for TT embedded in networks of the same orders but having different mean connectivities for: (i) ACoeff, (ii) BCoeff and (iii) Score. p-values of pairwise Mann-Whitney U tests are denoted by: ns - p > 0.05, * - 0.01< p <= 0.05, ** - 0.001 < p <= 0.01, *** - 0.0001 < p <= 0.001, **** - p <= 0.0001 **C)** Plots showing the dependence of change in the distribution of a metric with changing in-degree of a TT, for: (i) Score, (ii) ACoeff, and (iii) BCoeff. ACeoff and BCoeff are the coefficients corresponding to A and B terms, respectively, of the equation of plane given by linear multiple regression. Score is the r^2^ value of the linear regression model.

**Figure S10:**
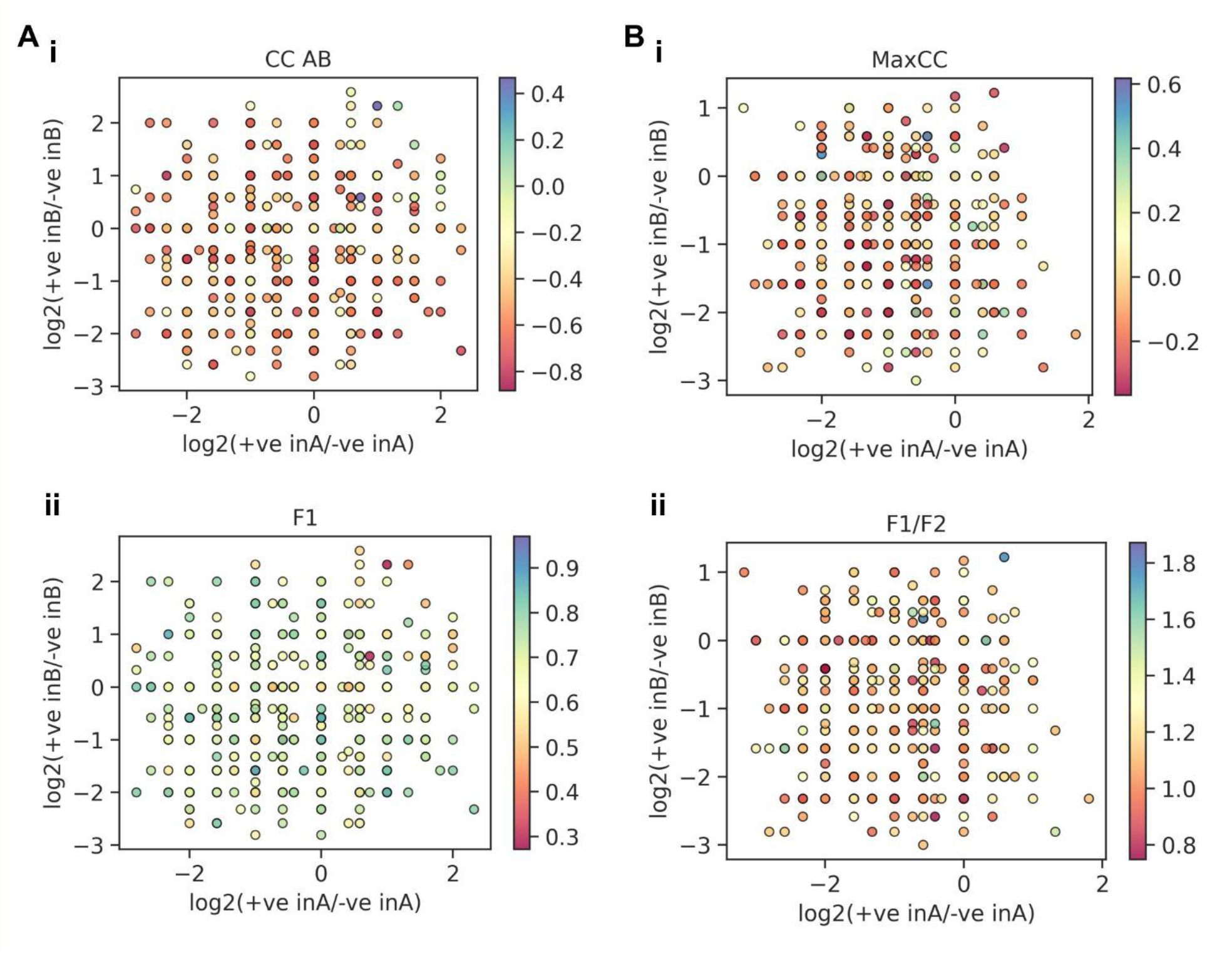
(A) Comparison of the variation of (i) CC AB and (ii) F1 values against the skew in positive to negative in-degrees of TS nodes. Each point is colored by their respective metric values as given by the color bar. (B) Comparison of the variation (i) MaxCC and (ii) F1/F2 values against the skew in positive to negative in-degrees of TT nodes. Each point is colored by their respective metric values as given by the color bar.

**Figure S11:**
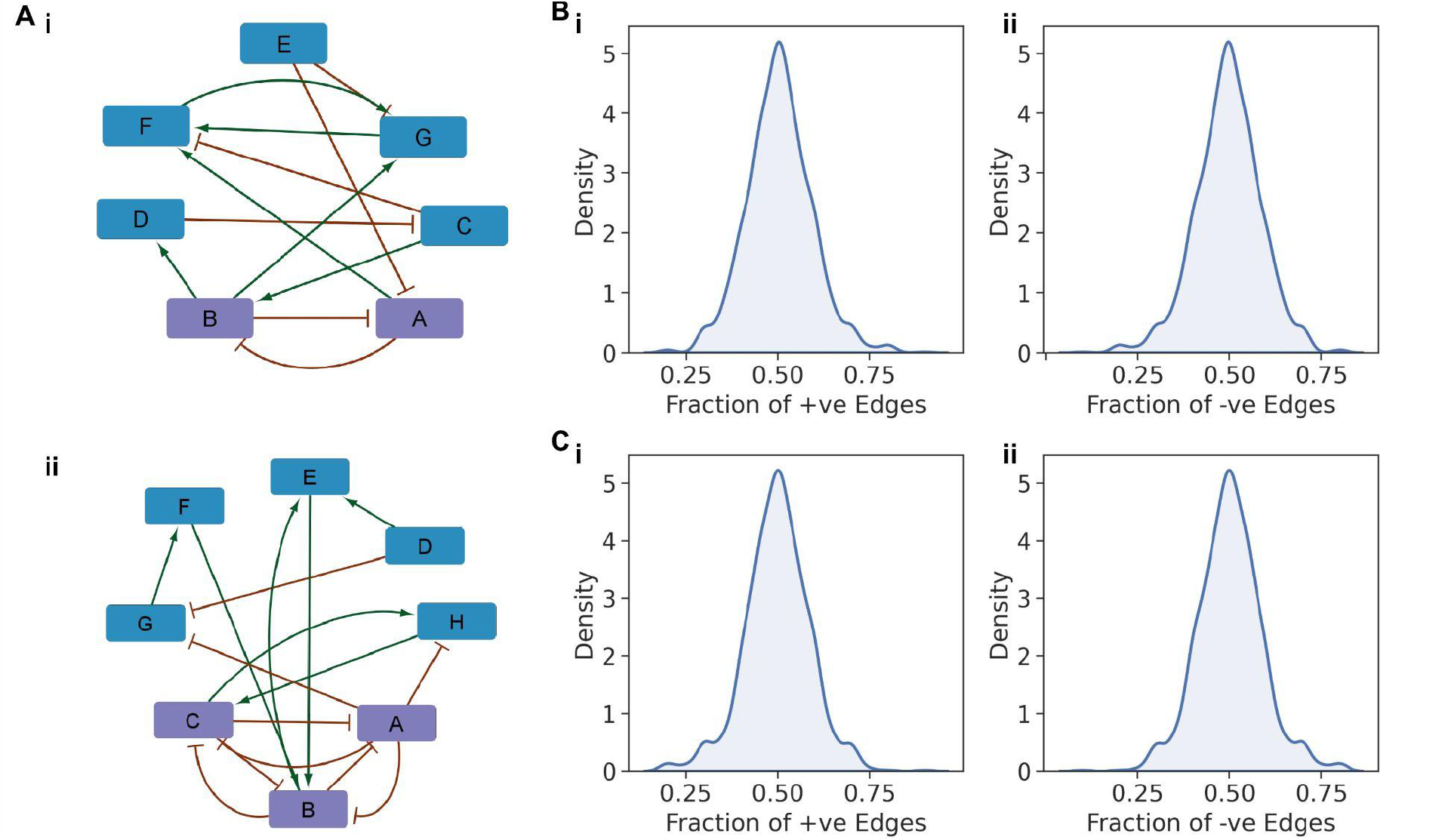
**A)** Representative network of 5N and E:2N i.e 10 edges with (i) TS embedded in it and (ii) TT embedded in it. TS and TT motifs are represented by purple colored nodes. **B)** Distribution of fraction of (i) positive and (ii) negative edges of the larger networks with embedded TS motifs. **C)** Distribution of fraction of (i) positive and (ii) negative edges of the larger networks with embedded TT motifs.

**Figure S12:**
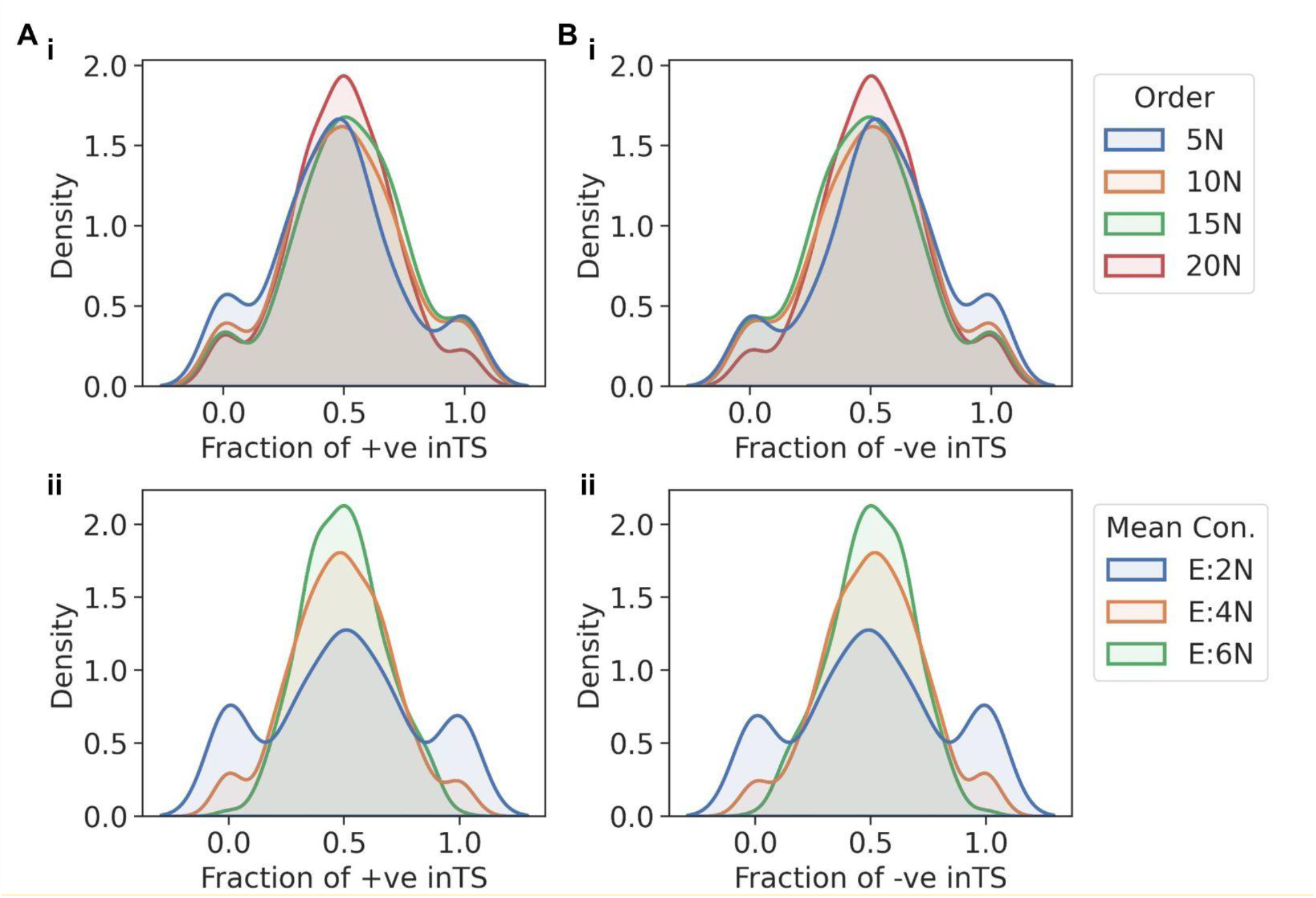
**A)** Distributions of fraction of positive in-degrees of TS motifs embedded in larger networks of i) differing network orders and ii) differing mean connectivities of networks. **B)** Distributions of fraction of negative in-degrees of TS motifs embedded in larger networks of i) differing network orders and ii) differing mean connectivities of networks.

**Figure S13:**
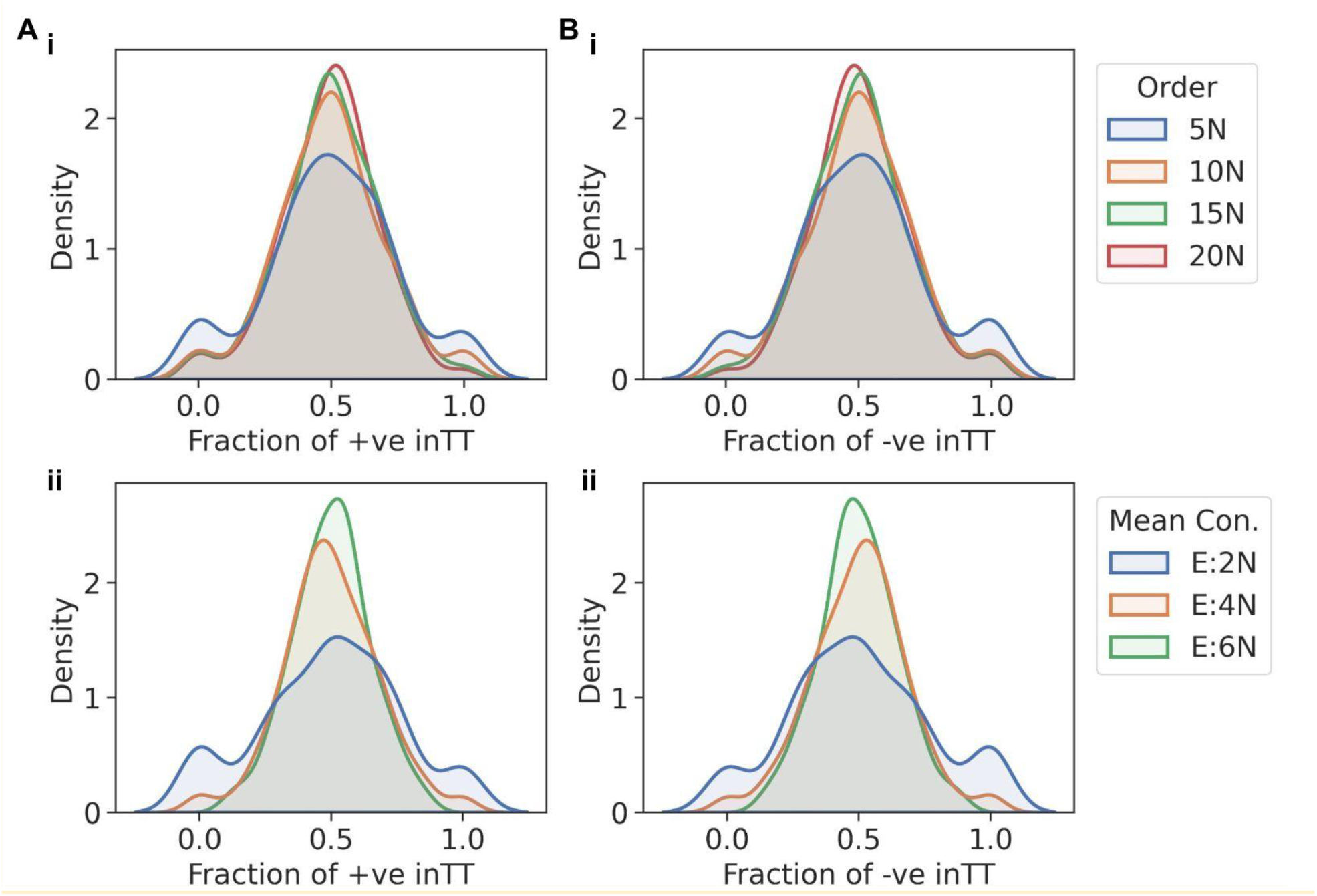
**A)** Distributions of fraction of positive in-degrees of TT motifs embedded in larger networks of i) differing network orders and ii) differing mean connectivities of networks. **B)** Distributions of fraction of negative in-degrees of TT motifs embedded in larger networks of i) differing network orders and ii) differing mean connectivities of networks.

